# Probabilistic quotient’s work & pharmacokinetics’ contribution: countering size effect in metabolic time series measurements

**DOI:** 10.1101/2022.01.17.476591

**Authors:** Mathias Gotsmy, Julia Brunmair, Christoph Büschl, Christopher Gerner, Jürgen Zanghellini

## Abstract

Metabolomic time course analyses of biofluids are highly relevant for clinical diagnostics. However, many sampling methods suffer from unknown sample sizes commonly known as size effects. This prevents absolute quantification of biomarkers. Recently, several mathematical post acquisition normalization methods have been developed to overcome these problems either by exploiting already known pharmacokinetic information or by statistical means.

Here we present an improved normalization method, MIX, that combines the advantages of both approaches. It couples two normalization terms, one based on a pharmacokinetic model (PKM) and the other representing a popular statistical approach, probabilistic quotient normalization (PQN), in a single model.

To test the performance of MIX, we generated synthetic data closely resembling real finger sweat metabolome measurements. We show that MIX normalization successfully tackles key weaknesses of the individual strategies: it (i) reduces the risk of overfitting with PKM, and (ii) contrary to PQN, it allows to compute sample volumes. Finally, we validate MIX by using real finger sweat as well as blood plasma metabolome data and demonstrate that MIX allows to better and more robustly correct for size effects.

In conclusion, the MIX method improves the reliability and robustness of quantitative biomarker detection in finger sweat and other biofluids, paving the way for biomarker discovery and hypothesis generation from metabolomic time course data.

## 1 Introduction

In recent years, the analysis of the sweat metabolome has received increased attention from several fields of study [1–3]. For example, sweat has been in the focus of forensic scientists since it is possible to analyze metabolomic profiles of finger prints that have been found (e.g. at a crime scene) [4]. Also, drug testing can easily be performed on sweat samples. One advantage of this method is to not only identify already illegal substances but their metabolic degradation products as well, thereby allowing to distinguish between drug consumption and mere contact [1]. Another application of sweat metabolomics is in diagnostics for personalized medicine, where the focus is put on discerning metabolic states of the body and trying to optimize nutrition and treatment based upon information of biomarkers in sweat [5–7].

Sweat metabolomics offers several technical advantages. Firstly, sweat is a rich source of biomolecules and thus offers great potential for biomarker discovery [8, 9]. Secondly, sweat sampling is easy compared to sampling of other biofluids (e.g. blood or urine). Moreover, it is non-invasive and can in principle be rapidly repeated.

Several sampling methods have been developed [2, 3, 9, 10]. However, most of them work in a very similar manner: a water absorbing material is put onto the skin’s surface to collect sweat for some (short) time. Sweat metabolites are subsequently extracted from this material and analyzed [3, 10]. Methods differ, however, in if and how they induce sweating. Some methods induce increased sweating by physical exercise [9] or chemical stimulation [2], whereas in other studies no sweat induction is performed and the natural sweat rate is sufficient for metabolomic analysis [3, 11].

Regardless of the exact sampling method, most of the above mentioned studies suffer from one major drawback. The sweat flux is highly variable, depending not only on interindividual differences, but also on body location, temperature, humidity, exercise and further factors that may change multiple times over the course of one day [12, 13]. For example, even with conservative estimates a variability of sweat flux *q*_sweat_ on the finger tips between 0.05 and 1 mg cm^-2^ min^-1^ needs to be accounted for [13–16]. This is a major challenge for comparative or quantitative studies, which has been acknowledged by many, e.g. [1, 4, 8, 17–19], however only actively approached by few – most notably [9].

The key problem is associated to the fact that often one is interested in the true metabolite concentrations, **C** ∈ ℝ^*n*_*metabolites*_^, of *n*_metabolites_ metabolites, which is obscured by an unknown and time-dependent sweat flux. Thus, the measured metabolites’ intensities are not proportional to **C** but to the metabolite mass vector, 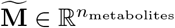,

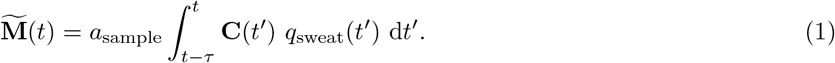

Here *a*_sample_ and *τ* denote the surface area of skin that is sampled, and the time it takes to collect one sample, respectively. We emphasize that throughout the manuscript the mass of a metabolite is defined as the measured abundance of the metabolite in a measured sample, and neither as the molar mass or mass to charge ratio. Moreover, we acknowledge that without a calibration curve the measured abundances have an arbitrary peak-area unit and are thus strictly neither absolute masses nor concentrations. The proportionality constant that scales measured intensities to mass units is determined by the calibration curve. The proper calibration curve is not further discussed here but assumed to be linear and available when applicable.

Metabolic concentration shifts happen in the span of double-digit minutes to hours, whereas sampling times are usually low single-digit minutes, therefore it is possible to assume that **C** changes little over the integration time τ [20]. Thus (1) simplifies to

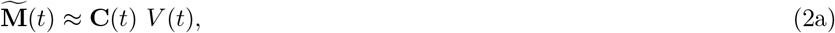

with an unknown sweat volume during sampling

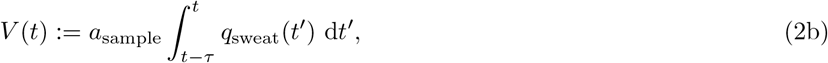

and the problems reads: given 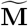, how can we compute **C** if we don’t know *V*?

The need to calculate absolute metabolite concentrations from small biological samples of unknown volume is not unique to sweat metabolomics, but known throughout untargeted metabolomics. The problem is commonly referred to as size effects [21]. For the sake of consistency with previous publications on this topic, we will use the term “size effects” throughout this publication. We emphasize that in this context it specifically refers to perceived differences in measured abundances due to changing sample volumes and/or dilutions and not to effects of different numbers of measurements per sample also referred to as sample size effects [22].

Three strategies have been developed to tackle size effects:

### Direct Sweat Volume Measurement

Measuring *V*, for instance via microfluidics [9, 23, 24], is the most straight forward method to solve (2) and typically very accurate with minimally required volumes in the range of ~ 5 to 100 μL [9, 23, 24]. However, in case of sweat sampling it may take quite some time, large sample areas or increased (i.e. induced) sweating to collect enough sweat for robust volume quantification. Another alternative is the volume estimation via paired standards [25], however, such method increases complexity of sample preparation. Either option would impede fast and easy sample collection and analysis.

### Indirect Sweat Volume Computation

If the chemical kinetics of targeted metabolite concentrations are known, then kinetic parameters and the sweat volume at each time point can be simultaneously determined by fitting the measured mass vector to Equation 2. Recently, we used this strategy to computationally resolve not only sample volumes in the nL to single digit μL-range but also accurately quantify personalized metabolic response patterns upon caffeine ingestion [20]. Albeit feasible for determination of individual differences with knowledge of reaction kinetics, this method quickly becomes unconstrained when too little prior information is available. Therefore, it is not suited for the discovery of unknown reaction kinetics. Moreover, this method requires several sampling time points to allow modeling the kinetics of different metabolites thereby decreasing simplicity of sampling.

### Statistical Normalization

With this approach the aim is to normalize the mass vector by the apparent mass of a marker that scales proportionally to the sample volume, so that the ratio becomes (at least approximately) independent of the sample volume. Various strategies have been developed for untargeted metabolomics; for example, normalization by total measured signal [26], and singular value decomposition-based normalization [27]. However, one of the best performing methods–referred to as probabilistic quotient normalization (PQN) – simply assumes that the median of the ratio of two apparent mass vectors is proportional to the sample volume [21, 28–30]. Although PQN does not allow one to compute sample volumes *per se,* it enables one to assess differential changes [28].

In this study we explore the performance of three different normalization methods on synthetic data. We illustrate the disadvantages of two previously published methods only focusing on either targeted or untargeted metabolites, respectively. A third normalization method is developed by combining both strategies in a single MIX model. We show that MIX significantly outperforms its preceding normalization methods. To validate the results we use MIX to characterize caffeine metabolization measured in the finger sweat as well as diphenhydramine metabolitzation measured in blood plasma.

## 2 Theory

### 2.1 Probabilistic Quotient Normalization

#### Definition

Probabilistic quotient normalization (PQN) assumes that for a large, untargeted set of metabolites the median metabolite concentration fold change between two samples (e.g. two measured time points, *t_r_* and *t_s_*) is approximately 1,

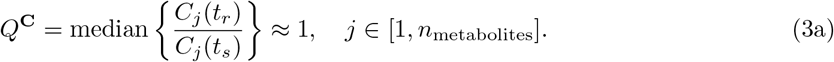

Consequently, fold changes calculated from 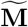 instead of **C** are proportional to the ratio of *V*,

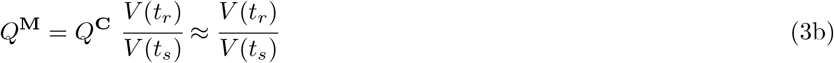

with

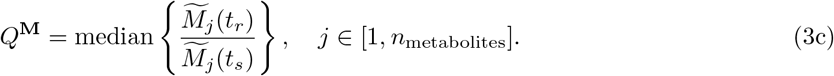

In order to minimize the influence of experimental errors

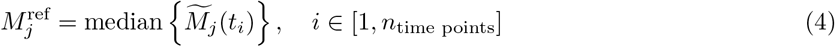

often replaces the dedicated sample in 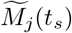 in the denominator of Equation 3c [28]. Therefore, the normalization quotient by PQN is calculated as

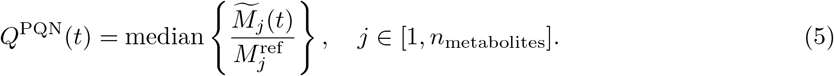

*Q*^PQN^ is a relative measure and distributes around 1. In analogy to Equation 3b, we define its relation to the (sweat) volume *V*^PQN^ as

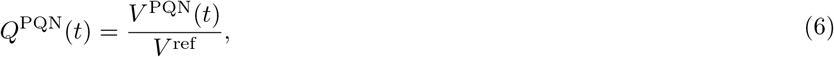

where *V*^ref^ denotes some unknown, time-independent reference (sweat) volume. Note that with real data only *Q*^PQN^(*t*) values can be calculated, but *V*^PQN^(*t*) as well as *V*^ref^ remain unknown.

#### Discussion

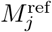 can be defined in different ways depending on the underlying data. However, the choice of of reference is usually not critical to the outcome of PQN [28]. As no control or blank measurements are available and the the abundances of metabolites can range several orders of magnitudes, in this study a metabolite-wise median reference was used for *Q*^PQN^ calculation. Moreover, PQN might be sensitive to missing values, however, in this study we only focused on (real and synthetic) data sets where 100% of values were present.

The biggest advantage of PQN is that no calibration curves and prior knowledge about changes over time of measured metabolites are required. Moreover, PQN is independent from the number of sample points measured in a time series. However, its major drawback is that the normalization quotient is not an absolute quantification and only shows relative changes. I.e. it does not quantify *V* as given in Equation 2 directly with an absolute value, but instead normalizes relative abundances between samples and time points. Another critical assumption is that sweat metabolite concentrations need to be – on average – constant over the sampled time series. Whereas this is reasonable to assume for the sweat of healthy humans [20], one has to take care when investigating disease states (for example cystic fibrosis, which is known to alter the sweat’s composition [31]).

### 2.2 Pharmacokinetic Normalization

#### Definition

In the pharmacokinetic model (PKM) we assume that we know at least the functional dependence, i.e. the pharmacokinetics, but not necessarily the value of the *k* (pharmaco-)kinetic parameters θ ∈ ℝ^*k*^ for 2 ≤ ℓ ≤ *n*_metabolites_ metabolites. Without loss of generality we (re-)sort 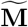 such that the first ℓ elements (collected in the vector 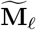) correspond to metabolites with known pharmacokinetic dependence, while the remaining *n*_metabolites_ – ℓ elements (collected in the vector 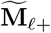) correspond to metabolites with unknown kinetics. Then Equation 2 takes the form of

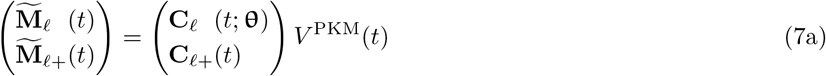

with physically meaningful bounds;

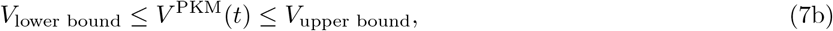

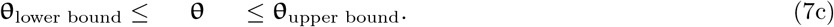

*V*^PKM^(*t*) as well as θ can be obtained by parametric fitting of 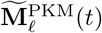. Note that this allows not only to compute absolute values of 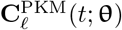 but – with *V*^PKM^(*t*) – also of all other concentrations via 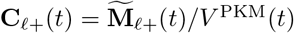.

As *V*^PKM^(*t_i_*) may be different at every time step *t_i_*, we need to know the (pharmaco-)kinetics of at least two metabolites, otherwise the number of parameters is larger than the number of data points.

#### Discussion

The biggest advantage of this method is that it can implicitly estimate absolute values of *V* without the need of direct measurements. Therefore, sweat volumes can become smaller than the minimum required in volumetric methods and shorter sampling times also become possible. A drawback of this method is the fact that it is only feasible if one has prior knowledge on relevant pharmacological parameters (i.e. ingested dose of metabolites of interest, volume of distribution, body mass of specimen, range of expected kinetic constants), which is limiting the approach to studies where at least two metabolites together with their pharmacokinetics are well known. Moreover, calibration curves of metabolites of interest, and sufficiently many samples in a time series are required for robustly fitting the equation system. In a previously performed sensitivity analysis, an increase in the quality of fit was observed as the number of samples increased from 15 to 20 time points per measured time series [20].

### 2.3 Mixed Normalization

#### Definition

The mixed normalization model (MIX) is a combination of PQN and PKM. It is designed to incorporate robust statistics of untargeted metabolomics via its PQN term as well as an absolute estimation of *V* via its PKM term.

Optimal parameters of MIX are found via optimization of two equations,

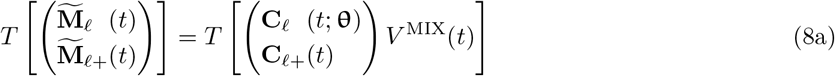

and

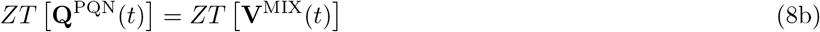

where additional transformations *T* (PKM and PQN term) and scaling *Z* (PQN term) can be applied to account for random and systematic errors (Section 3.1.1) and *V*^MIX^(*t*) and θ are constrained between physically meaningful bounds,

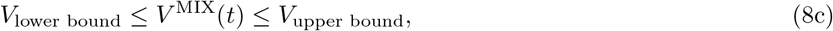

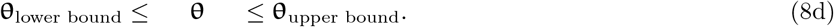

E.g. bounds for *V* can be calculated by Equation 2a and minimal and maximal sweat rates from literature.

#### Discussion

We hypothesize that MIX model can combine the advantages of PQN and PKM normalization models. Moreover, we believe that MIX inherits the statistical robustness of PQN while simultaneously estimating absolute values as fitted by PKM. Several prerequisites are necessary for normalization with PKM or MIX. However, if they are fulfilled, the improved goodness of normalization by using MIX instead of PKM usually does not come with an additional price as in many metabolomics studies targeted and untargeted metabolites are measured in combination and thus all additional data required by MIX is already available.

## 3 Methods

### 3.1 Implementation

A generalized version of PKM and MIX (where an arbitrary number of independent metabolite kinetics can be modeled) was implemented as a Python class. As input it requires the number of metabolites used for kinetic modeling (ℓ), a vector of time points as well as the measured mass data (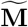, matrix with time points in the rows and metabolites in the columns). MIX additionally takes a *Q*^PQN^ = [*Q*^PQN^(*t*_1_),…, *Q*^PQN^(*t*_*n*_time points__)]^T^ vector (calculated with the PQN method from all metabolites, *n*_metabolites_) for all time points of a time series. Upon optimization (carried out with `self.optimize_monte_carlo,` which is a wrapper for SciPy’s `optimize.curve_fit` [32]) the kinetic constants and sweat volumes are optimized to the measured data by minimizing the functions listed in Equations 9b and 9c for PKM and MIX respectively:

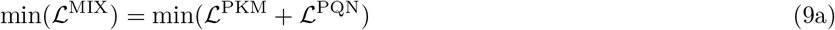

where

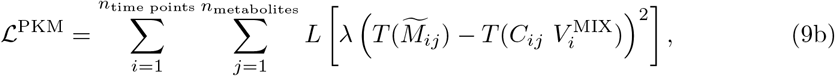

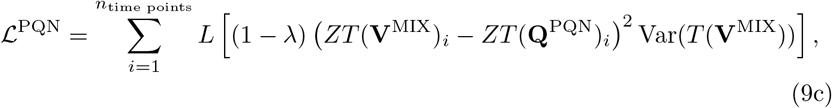

Var(**V**) is the variance of **V** (which is the vector of estimated *V* over all time points), *T* is a transformation function, *Z* is a scaling function, and *L* is the loss function. The key difference between PKM and MIX is that the fitted *V* in MIX are biased towards relative abundances as calculated by PQN. An important additional hyperparameter of the MIX model is λ, which weights the error residuals of 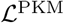 and 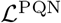. Its calculation is discussed in Section 3.1.1. If λ =1, the MIX model simplifies again to a pure PKM model.

To summarize, an overview of the differences of PKM and MIX model is given in Supplementary Table S1 and a flow chart of data processing for MIX normalization is given in Supplementary Figure 1.

#### 3.1.1 Hyperparameters

Several hyperparameters can be set for the PKM and MIX Python classes.

##### Kinetic Function

Firstly, it is possible to choose the kinetic function used to calculate **C**. In this study we focused on a modified Bateman function *F*(*t*) with 5 kinetic parameters (*k_a_*, *k_e_, c*_0_, *lag, d*):

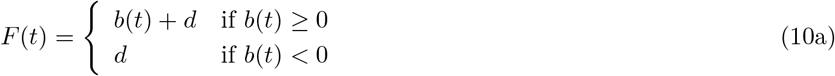

with

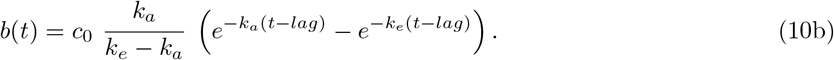

This function was designed to be flexible and able to represent several different metabolite consumption and production kinetics as exemplified by Figure 2. Intuitively, *k_a_* and *k_e_* correspond to kinetic constants of absorption and elimination of a metabolite of interest with the unit h^-1^. *c*_0_ is the total amount of a metabolite absorbed over the volume of distribution with the unit mol L^-1^. Additionally to these parameters which are also part of the classical Batman function [33], we here introduce *lag* and *d.* The *lag* term with the unit h shifts the function along the X-axis, intuitively defining the starting time point of absorption of a metabolite of interest, whereas the *d* term with the unit mol L^-1^ shifts the function along the Y-axis.

**Figure 1.**
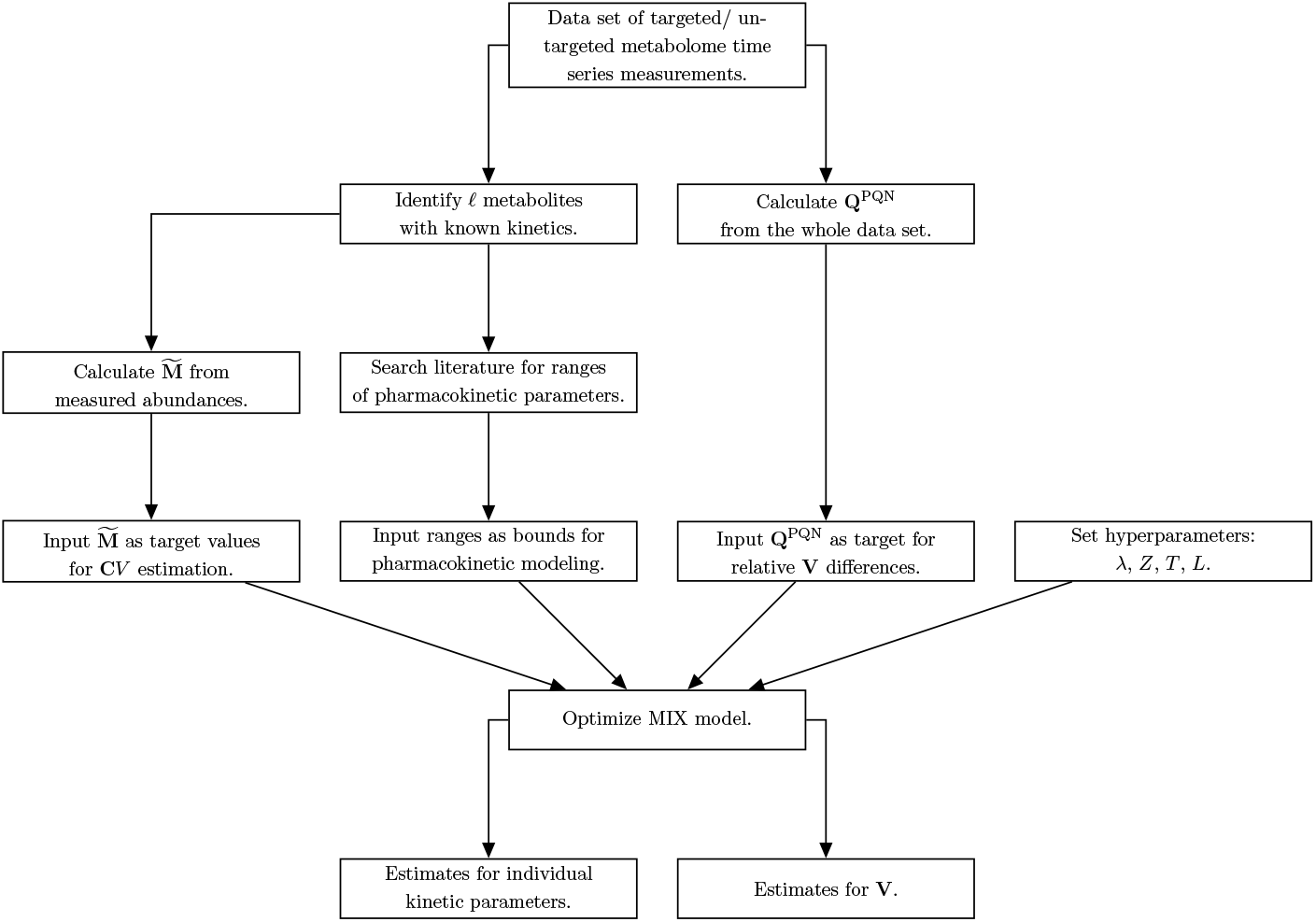
Flow chart for data processing for MIX normalization.

**Figure 2.**
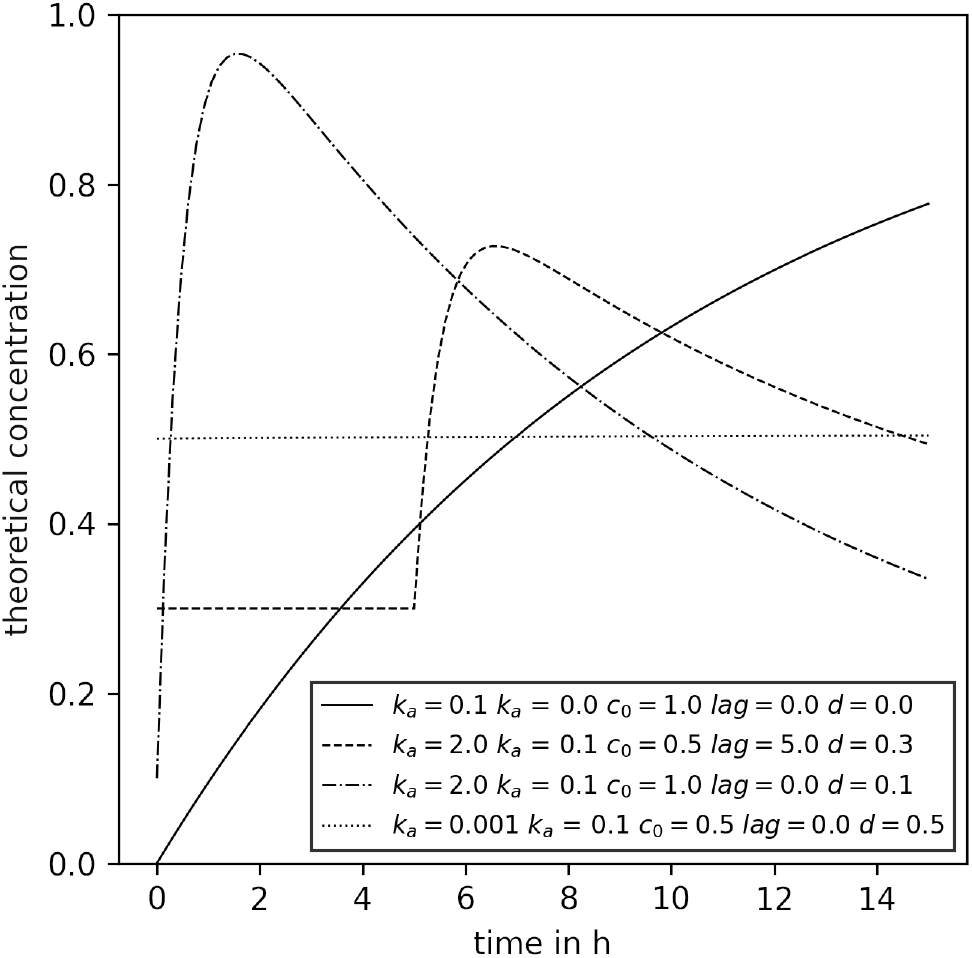
Examples of concentration time series that can be modeled with the modified Bateman equation used. The legend shows the kinetic parameters used to create the respective curves. All parameters are within the bounds that were used for kinetic parameter fitting.

##### Loss Function

*L. L* calculates the loss value after estimation of the error residuals of the model (Equation 9). It can be set via `self.set_loss_function` to either `cauchy_loss` or `max_cauchy_loss` (or `max_linear_loss`). In both cases the loss is calculated as a Cauchy distribution of error residuals according to SciPy [32]. The difference, however, is that `cauchy_loss` only uses the absolute error residuals, whereas `max_cauchy_loss` uses the maximum of relative and absolute error residuals (thus the word max is expressed in its name). The reason for its addition was that a good performance has been achieved in a previous study [20]. In this study we used the `max_cauchy_loss` loss function for PKM models and `cauchy_loss` for MIX models. The choice of *L* is intertwined with the choice of *T* which becomes clear in the following paragraph.

##### Transformation Function

*T. T* transforms the measured data 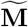 as well as the calculated *Q*^PQN^, **C***V*, and **V** before calculation of the loss (Equation 9). Two different transformations, none and `log10`, can be set during initialization with the argument `trans_fun`. As originally reported [20] no transformation was done for PKM (i.e. `trans_fun=‘none’`),

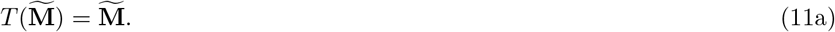

For MIX models, however, a log-transform was performed (i.e. `trans_fun=‘ log10’`),

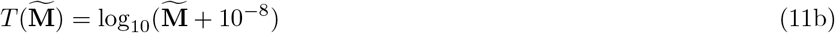

as the error on measured data is considered multiplicative [34] and the sweat volume log-normally distributed (Supplementary Figure S1). To avoid problems with concentrations of the size 0 a small number (i.e. the size of optimizer precision [32]) is added.

In a sensitivity analysis study, we tested the quality of normalization of MIX with different *L* and *T* hyperparameters and concluded that a combination of `cauchy_loss` for *L* and `log10` for *T* performed best (Supplementary Figure S2C, D).

##### Scaling Function

*Z. Z* describes a scaling function performed on *T*(*Q*^PQN^) and *T*(**V**). Scaling is performed to correct for noisy data (see Results Section 4.2.1). Two strategies can be set with the `scale_fun` argument during initialization of the MIX model class, `standard` or `mean`. In this study, all MIX models employ standard scaling, i. e.

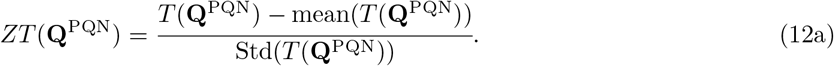

We additionally implemented `mean` scaling which differs depending on the choice of *T* with

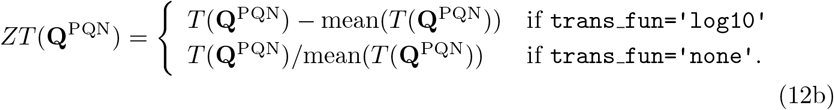

##### Optimization Strategy

The optimization of both, PKM and MIX models, is done with a Monte Carlo strategy where the initial parameters sampled randomly from an uniform distribution between their bounds. Performing a sensitivity analysis, we previously showed that this method is preferable to a single fitting procedure [20]. In this study the number of Monte Carlo replicates for model fitting was set to 100.

##### Weighting of MIX Loss Terms

A weighting constant for every measured data point can be used by the model. In a sensitivity analysis study we found that the choice of λ is not critical to the quality of normalization as long as it is not extremely tilted to one side (i.e. λ close to 0 or 1, Supplementary Figure S2A, B). Thus we propose a method where the loss terms are weighted by the number of data points fitted for each of both loss terms, but not by the number of metabolites used in the calculation of each term (Supplementary Equations S1). For such a method the solution for λ is given by Equation 13.

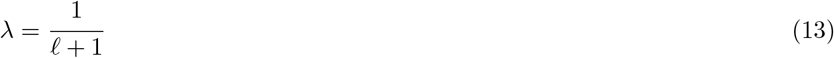

#### 3.1.2 Full and Minimal Models

In this study we differentiate between full and minimal models. With full models we refer to pharmacokinetic normalization models (PKM or MIX) where all metabolites of a given data set are used for the pharmacokinetic normalization. This means that, for example, if *n*_metabolites_ = 20 all 20 metabolites were modeled with the modified Bateman function and thus in Equations 7a and 8a, ℓ = *n*_metabolites_ and 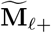 is an empty vector. On the other hand, minimal models are models where only the few, known, better constrained metabolites were modeled with a kinetic function. This means that the information used for PKM_minimal_ does not change upon addition of synthetic metabolites. Therefore, its goodness of fit measure should stay constant within statistical variability upon change of *n*_metabolites_. This behaviour was used to verify if the simulations worked as intended and no biases in the random number generation exist. On the other hand MIX_minimal_ model still gained information from the increase of *n*_metabolites_ as the PQN part of this model was calculated with all *n*_metabolites_. Therefore, changes in the goodness of fit measures for MIX_minimal_ are expected. We emphasize that the definition of full and minimal models is specific to this particular study. Here we explicitly set ℓ = 4, which originates from previous work where 4 targeted metabolites (caffeine, paraxanthine, theobromine, theophylline) with known kinetics were measured [20].

### 3.2 Synthetic Data Creation

Three different types of synthetic data sets were investigated. The first two types of data sets (sampled from kinetics, Section 3.2.1 and sampled from means and standard deviations, Section 3.2.2) test the behaviour of normalization models in extreme cases (either all metabolites describable by pharmacokinetics or all metabolites completely random). Finally, the third type of data set (sampled from real data, Section 3.2.3) aims to replicate measured finger sweat data as close a possible. In sum the performance of normalization methods on all three types of data sets can show how they behave in different situations with different amounts of describable data.

In all three cases data creation started with a simple toy model closely resembling the concentration time series of caffeine and its degradation products (paraxanthine, theobromine, and theophylline) in the finger sweat as described elsewhere [20]. The respective parameters are listed in Supplementary Table S2. With them the concentration of metabolites #1 to #4 were calculated for 20 time points (between 0 and 15 h in equidistant intervals, Figure 3). Subsequently, new synthetic metabolite concentration time series were sampled and appended to the toy model (i.e. to the concentration vector, **C**(*t*)). Three different synthetic data sampling strategies were tested and their specific details are explained in the following sections. Next, sweat volumes (*V*) were sampled from a log-normal distribution truncated at (0.05 ≤ *V* ≤ 4 μL) closely resembling the distribution of sweat volumes estimated in our previous publication [20], Supplementary Figure S1. Finally, an experimental error (ϵ) was sampled for every metabolite and time point from a normal distribution with a coefficient of variation of 20% and the synthetic data was calculated as

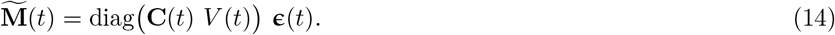

For every tested condition 100 synthetic data replicates were generated and the normalization models were fitted.

**Figure 3.**
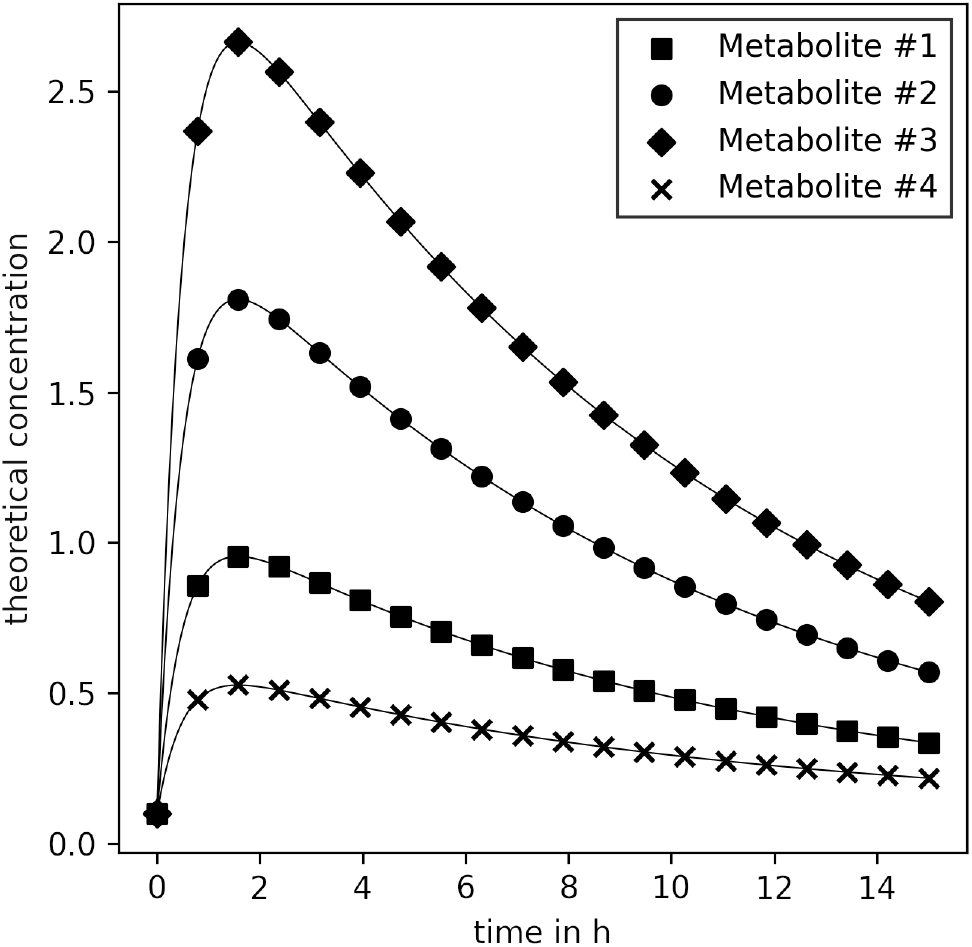
C for the first four metabolites of the synthetic data. Kinetic parameters used for calculation are listed in Supplementary Table S2.

#### 3.2.1 Sampled Kinetics

In simulation v1, data was generated by sampling kinetic parameters for new metabolites from an uniform distribution. The distribution was constrained by the same bounds also used for the PKM and MIX model fitting: (0, 0, 0, 0)^T^ ≤ (*k_a_, k_e_, c*_0_, *lag, d*)^T^ ≤ (3, 3, 5, 15, 3)^T^. Subsequently the concentration time series of the synthetic metabolites were calculated according to the modified Bateman function (Equation 10).

#### 3.2.2 Sampled Mean and Standard Deviation

Means and standard deviations of the concentration time series of metabolites were calculated from untargeted real finger sweat data (for details see Section 3.4). The probability density function of both can be described by a log-normal distribution (Supplementary Figure S3). For the data generation of simulation v2, per added metabolite one mean and one standard deviation were sampled from the fitted distribution and used as an input for another log-normal distribution from which a random concentration time series was subsequently sampled. This results in synthetic concentration values that behave randomly and, therefore, cannot be easily described by our pharmacokinetic models.

#### 3.2.3 Sampled from Real Data

To get an even better approximation to real data, in simulation v3 concentration time series were directly sampled from untargeted real finger sweat data (for details see Section 3.4). To do so, the untargeted metabolite 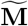 time series data set was normalized with PQN. As the number of metabolites in this data set was comparably large (*n*_metabolites_ = 3446) we could assume that the relative error (or rRMSE, for more explanation see Section 4.1.1) was negligibly small. The resulting values are strictly speaking fractions of concentrations. However, this does not affect the results as these values are anyways considered untargeted (i.e. no calibration curve exists) and thus relative. Therefore, the PQ normalized data set could be used as ground truth for concentration time series sampling. Subsequently, a subset of the original ground truth data was sampled for synthetic data generation.

#### 3.2.4 Sampling of Noisy Data

We investigated the influence of background (i.e. noisy) signal on the performance on **Q**^PQN^ (and scaled and transformed variants thereof). To simulate such an environment we used data sampled from real data (Section 3.2.3), and applied *V* only to a fraction of the **C** vector,

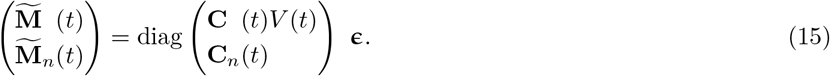

The noise fraction is given by the number of elements of 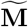 and 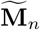 vectors,

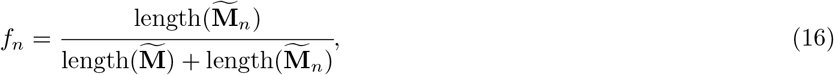

where subscript *n* in 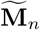, **C**_*n*_, and *f_n_* denotes them as part of the noise.

Simulations were carried out for 20 equidistant noise fractions between 0 ≤ *f_n_* ≤ 0.95 with *n*_metabolites_ = 100 and *n*_time points_ = 20 for 100 replicates. The error residuals of mean and standard scaled **Q**^PQN^ are calculated as

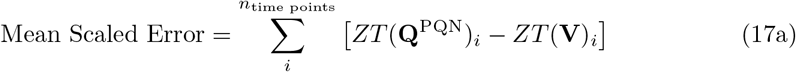

with *Z* defined as in Equation 12b and

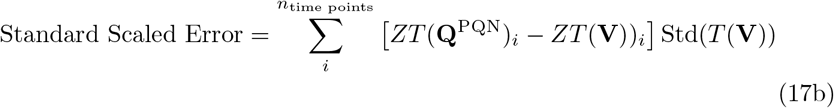

with *Z* defined as in Equation 12a. For both cases *T* is defined as the logarithm (Equation 11b). We point out that the multiplication with Std(*T*(**V**)) for the standard scaled error is important to make the results comparable, as otherwise the error would be biased towards the method with smaller scaled standard deviation regardless of the performance of the scaling.

### 3.3 Normalization Model Optimization

Normalizing for the sweat volume by fitting kinetics through the measured values only has a clear advantage over PQN if it is possible to infer absolute sweat volumes and concentration data. In order to be able to do that, some information about the kinetics and the starting concentrations of metabolites of interest need to be known. For example, when modeling the caffeine network in our previous publication [20] we knew that the *lag* parameter of all metabolites was 0 and that the total amount of caffeine ingested (which corresponds to *c*_0_) was 200 mg. Moreover, we knew that caffeine and its metabolites are not synthesized by humans and implemented the same strategy into our toy model (corresponding to *d*). As the toy model was designed to resemble such a metabolism we translated these information to the current study. Therefore, we assumed that the first 4 metabolites in our toy model had known *c*_0_, *lag*, and *d* parameters. For their corresponding *k_a_* and *k_e_* and the parameters of all other metabolites the bounds were set to the same (0, 0, 0, 0)^T^ ≤ (*k_a_, k_e_, c*_0_, *lag, d*)^T^ ≤ (3, 3, 5, 15, 3)^T^ used in kinetic data generation. Figure 2 shows examples of concentration time series that can be described with the modified Bateman function and parameters within the fitting bounds.

### 3.4 Real Finger Sweat Metabolome Data

The real world finger sweat data was extracted from 37 time series measurements of Study C from ref. [20]. It was downloaded from MetaboLights (MTBLS2772 and MTBLS2776).

#### Preprocessing

The metabolome data set was split into two parts: targeted and untargeted. The targeted data (i.e. the mass time series data for caffeine, paraxanthine, theobromine, and theophylline) was directly adopted from the mathematical model developed by [35]. This data is available on GitHub (https://github.com/Gotsmy/finger_sweat).

For the untargeted metabolomics part, the raw data was converted to the mzML format with the msConvert tool of ProteoWizard (version 3.0.19228-a2fc6eda4) [36]. Subsequently, the untargeted detection of metabolites and compounds in the samples was carried out with MS-DIAL (version 4.70) [37]. A manual retention time correction was first applied with several compounds present in the majority (more than 90%) of the samples. These compounds were single chromatographic peaks with no isomeric compounds present at earlier or later retention times (*m/z* 697.755 at 5.57 min, m/z 564.359 at 5.10 min, *m/z* 520.330 at 4.85 min, *m/z* 476.307 at 4.58 min, *m/z* 415.253 at 4.28 min, *m/z* 371.227 at 3.95 min, *m/z* 327.201 at 3.56 min, *m/z* 283.175 at 3.13 min, *m/z* 239.149 at 3.63 min, *m/z* 166.080 at 1.69 min, *m/z* 159.113 at 1.19). After this, untargeted peak detection and automated alignment (after the manual alignment) were carried out with the following settings: Mass accuracy MS1 tolerance: 0.005 Da, Mass accuracy MS2 tolerance: 0.025 Da, Retention time begin: 0.5 min, Retention time end: 6 min, Execute retention time correction: yes, Minimum peak height: 1E5, Mass slice width: 0.01 Da, Smoothing method: Linear weighted moving average, Smoothing level: 3 scans, Minimum peak width: 5 scans, Alignment reference file: `C_D1_I_o_pos_ms1_1.mzML`, Retention time tolerance: 0.3 min, MS1 tolerance: 0.015 Da, Blank removal factor: 5 fold change). No blank-subtraction was carried out as the internal standard caffeine was spiked into each sample including the blanks. Peak abundances and meta-information were exported with the `Alignment results export` functionality.

Subsequently, we excluded isomers within a *m/z* difference of less than 0.001 Da and a retention time difference of less than 0.5 min. To further reduce features that are potentially background, features with retention times after 5.5 min as well as features with minimal sample abundances of < 5 × maximum blank abundance (except for the internal standard, caffeine-D9) were excluded from the data set. This was done on a time series-wise basis, thus the number of untargeted metabolites considered for normalization differs with a mean of 343 ± 152 for the 37 time series of interest.

#### Size Effect Normalization

In this finger sweat data set, time series of targeted as well as untargeted metabolomics are listed. The kinetics of the four targeted metabolites (caffeine, paraxanthine, theobromine, and theophylline) are known. A reaction network of the metabolites is shown in the top panel of Figure 4. Briefly, caffeine is first absorbed and then converted into three degradation metabolites. Additionally, all four metabolites are eliminated from the body. All kinetics can be described with first order mass action kinetics [38, 39].

**Figure 4.**
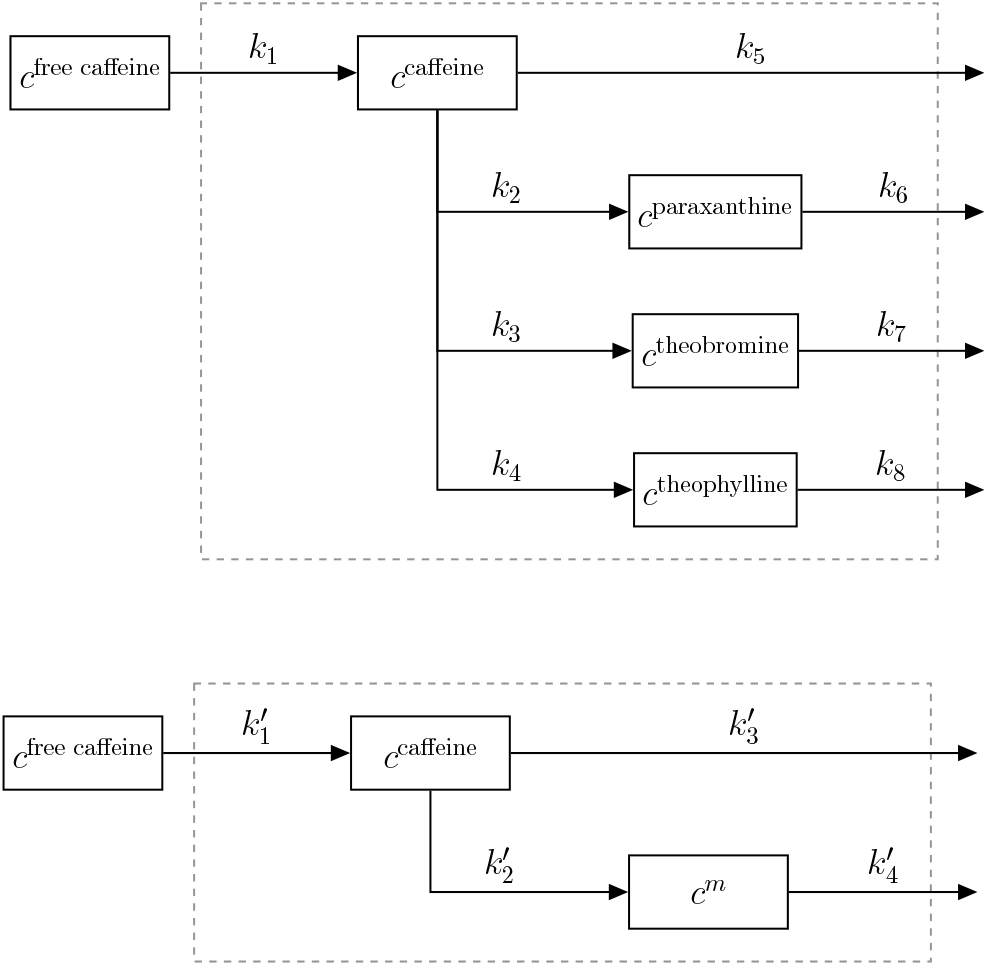
Full network (top panel) and subnetwork (bottom panel) of caffeine absorption, conversion to paraxanthine, theobromine, and theophylline and their elimination. The system boundary (dashed line) represents the human body. *m* ∈ {paraxanthine, theobromine, and theophylline}

In order to assess the performance of the sweat volume normalization methods the full network was split up into three subnetworks that all contained caffeine and one degradation metabolite each (Figure 4 bottom panel). The solution of the first order differential equations describing such network is given in Supplementary Equations S2a and S2b. Moreover, the 343 ± 152 untargeted metabolite time series were randomly split up into three (almost) equally sized batches and each batch was assigned to one subnetwork. All three networks were subsequently separately normalized with PKM_minimal_ and MIX_minimal_ methods with kinetic parameters that were adjusted to the specific reaction network (Figure 4 bottom panel). Subsequently, the kinetic constants 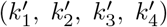 were estimated for 37 measured concentration time series. Fitting bounds were not changed in comparison to the original publication [20].

As all three subnetwork data sets originate from the same finger sweat measurements, the underlying kinetic constants should be exactly identical. As the kinetic constants of absorption 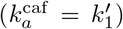 and elimination 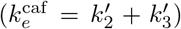 of caffeine are estimated in all three subnetworks we used their standard deviation to test the robustness of the tested normalization methods.

### 3.5 Real Blood Plasma Metabolome Data

In the study of Panitchpakdi et al. [40] the mass time series of the metabolome was measured in different body fluids after the uptake of diphenhydramine (DPH). Here, we focus on data measured in the blood plasma which includes the abundances of DPH (known kinetics, calibration curve, pharmacological constants) as well as three of its metabolization products (known kinetics) and the abundances of 13526 untargeted metabolites with unknown kinetics.

#### Preprocessing

The data of peak areas was downloaded from the GNPS platform [41]. To reduce the number of metabolites that are potentially background and/or noise in the data set, features with minimal sample abundances of < 5 × maximum blank abundance were excluded from the data set on a time series-wise basis. Thus, the number of untargeted metabolites considered for normalization differs with a mean of 1017 ± 114 for the 10 time series of interest.

#### Size Effect Normalization

We assume that the kinetics of four metabolites (DPH, N-desmethyl-DPH, DPH N-glucuronide, and DPH N-glucose) can be described by the modified Bateman (Equation 10). A reaction network of the metabolites is shown in Supplementary Figure S4. Briefly, DPH is first absorbed and then – with unknown intermediates – converted into three degradation metabolites, which are in turn metabolized further downstream or eliminated. *c*_0_ of DPH was calculated with pharmacological constants for bioavailability, volume of distribution, and dosage of DPH as reported in the original publication [40].

Analogously to the normalization performed on finger sweat data, the full network of four metabolites is split up into three subnetworks with only one, shared, targeted metabolite (DPH itself), one additional untargeted metabolite with known kinetic (either N-desmethyl-DPH, DPH N-glucuronide, or DPH N-glucose, Supplementary Figure S5) and one third of 1017 ± 114 untargeted metabolites with unknown kinetics. To ensure better convergence during fitting of the models, the 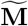 data was first scaled to values between 0 and 1 by dividing by its metabolite-wise maximum. This factor can be multiplied again as part of *c*_0_ after the normalization is done. Thereafter, PKM_minimal_ and MIX_minimal_ models were fitted onto the scaled 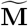 data (with ℓ =2) for all ten measured time series. The bounds of parameters were chosen so that previously reported estimates [40] are well within range: 0 ≤ *k* ≤ 5h^-1^ for 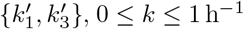 for 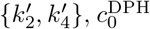 as reported in the original publication normalized by the maximum factor, 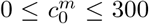 for *m* ∈ {N-desmethyl-DPH, DPH N-glucuronide, DPH N-glucose} and *lag* = *d* = 0 as well as 0.01 ≤ *V* ≤ 0.03 mL.

As all three subnetwork data sets originate from the same plasma time series measurements, the underlying kinetic constants of DPH should be exactly identical. As the kinetic constants of absorption 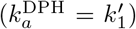 and elimination 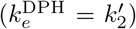 of DPH are estimated in all three subnetworks we used their standard deviation to test the robustness of PKM_minimal_ and MIX_minimal_.

### 3.6 Data Analysis

#### Goodness of Normalization

Two goodness of fit measures are calculated to analyze the performance of the tested methods. RMSE is the standard deviation of the residuals of a sampled sweat volume time series vector (**V**^true^) minus the fitted sweat volume vector (**V**^fit^), while rRMSE is the standard deviation of the ratio of sampled and fitted **V** vectors normalized by its mean. Intuitively, RMSE is a measure of how much absolute difference there is between the fit and a true value, rRMSE on the other hand gives an estimate on how good the fitted sweat volumes are relative to each other. A visual depiction of RMSE and rRMSE is shown in Supplementary Figure S6 and their exact definition is given in the equations in 3.3.

#### Statistical Analysis

The significant differences in the mean of goodness of fit measures were investigated by calculating *p* values with the non-parametric pairwise Wilcoxon signed-rank test [42] (SciPy’s `stats.wilcoxon` function [32]). Significance levels are indicated by *, **, and *** for p ≤ 0.05, 0.01, and 0.001 respectively.

## 4 Results

### 4.1 Comparison of PKM and MIX

#### 4.1.1 Synthetic Data Simulations

In order to test the performance of different normalization models we generated 100 synthetic data sets with three different methods (simulations v1, v2, v3) and five different *n*_metabolites_ (4, 10, 20, 40, 60) each, where the underlying **C**, *V*, and ϵ values were known. Simulations v1, v2, and v3 differ in the way how **C** was generated (kinetic, random, sampled from real data set, respectively). In order to quantify the normalization model performance, two measures of goodness of normalization were used for the analysis of the results: RMSE and rRMSE.

To visualize the obtained normalization performances we plotted the results for simulation v3 and *n*_metabolites_ = 60 in Figure 5 for three normalization models (from left to right column, PQN, PKM_minimal_, and MIX_minimal_). The top row shows the predicted log_10_(*C_j_*(*t_i_*; θ)/*C_j_*(0; θ)) (i.e. the concentration of each metabolite *j* at each time point *i* divided by its concentration at time 0) as a function of the true log_10_(*C_j_* (*t_i_*)/*C_j_* (0)) values. It illustrates the correlation of the relative abundances of one metabolite across all time points. Good correlations (i.e. high R^2^) as seen for PQN and MIX_minimal_ result in a low rRMSE measure. On the bottom row of Figure 5 the absolute values of predicted *V* are plotted as a function of the true *V*. There it becomes evident that good correlations of absolute values result in low RMSE measures.

**Figure 5.**
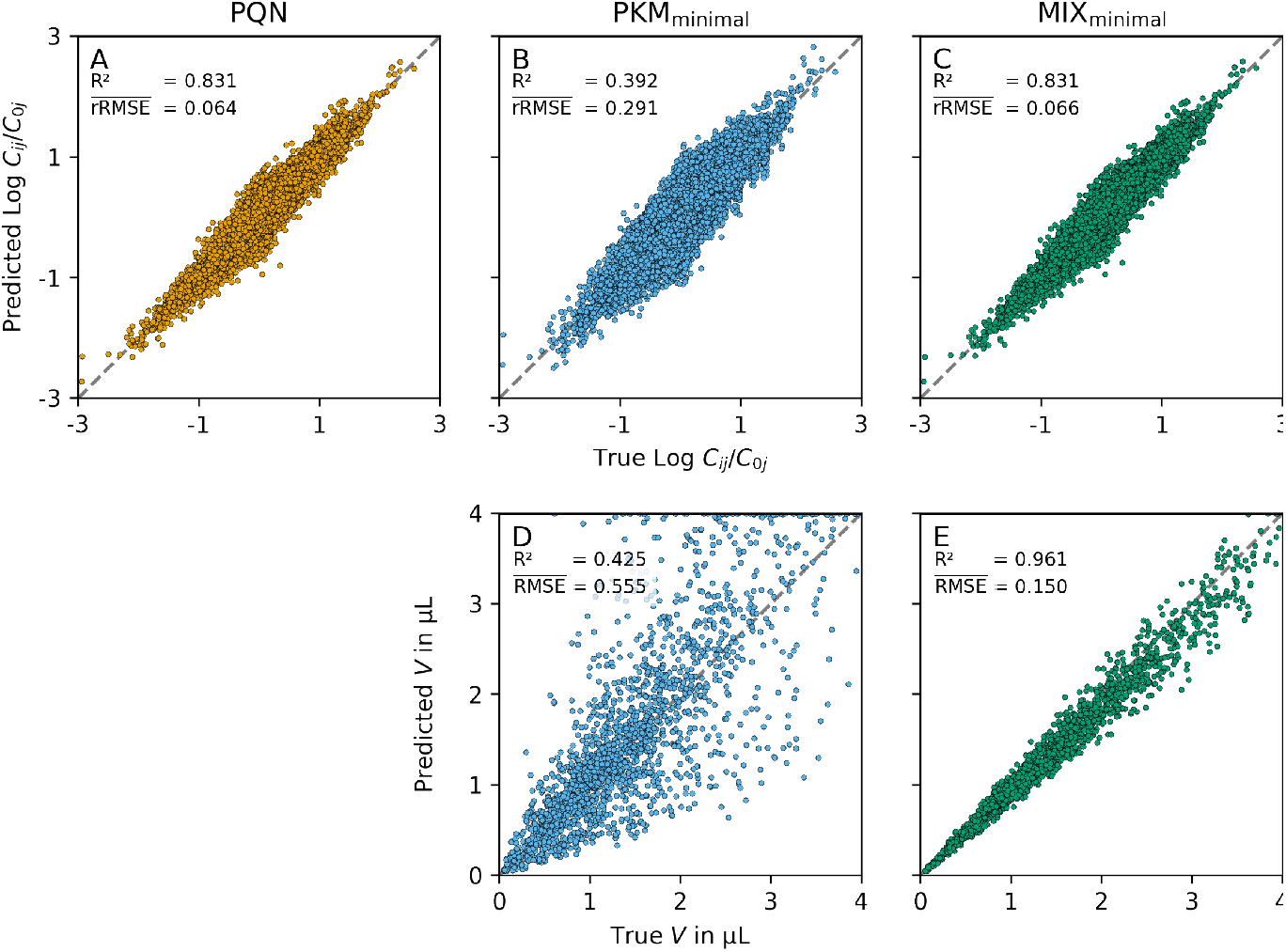
Relative and absolute normalization performance. In the top row the predicted log_10_(*C_j_* (*t_i_*; θ)/*C_j_* (0; θ)) (1 ∈ {*n*…, *n*_time points_}, *j* ∈ {1,…, *n*_metabolites_}) are plotted as a function of the true, underlying log_10_ (*C_j_* (*t_i_*)/*C_j_* (0)). The bottom row shows the predicted *V* as a function of the true, underlying **V**. The columns represent different normalization models (PQN, PKM_minimal_, and MIX_minimal_ from left to right). As no absolute **V** can be calculated from PQN the bottom left plot is omitted. To illustrate the effect of different RMSE and rRMSE sizes (which both are calculated from **V**), we show their mean over 100 replicates in comparison to the R^2^ values calculated from the points plotted. Intuitively rRMSE is a measure of good correlation on the top row whereas RMSE is a measured of good correlation on the bottom row (high R^2^, low rRMSE/RMSE respectively).

In the following sections we will focus on the size of RMSE and rRMSE respectively as they are both calculated from the predicted *V* directly. Note that for PQN no absolute *V* can be estimated and, therefore, no RMSE is calculated.

##### Influence of the Number of Metabolites

We tracked RMSE and rRMSE of normalization methods for different numbers of metabolites (*n*_metabolites_) to investigate how the methods behave with different amounts of available information. An overview of their goodness of normalization measures as a function of *n*_metabolites_ on sampled kinetic data (panels A, B), on completely random data (panels C, D) and on sampled subsets of real data (panels E, F) is given in Figure 6.

**Figure 6.**
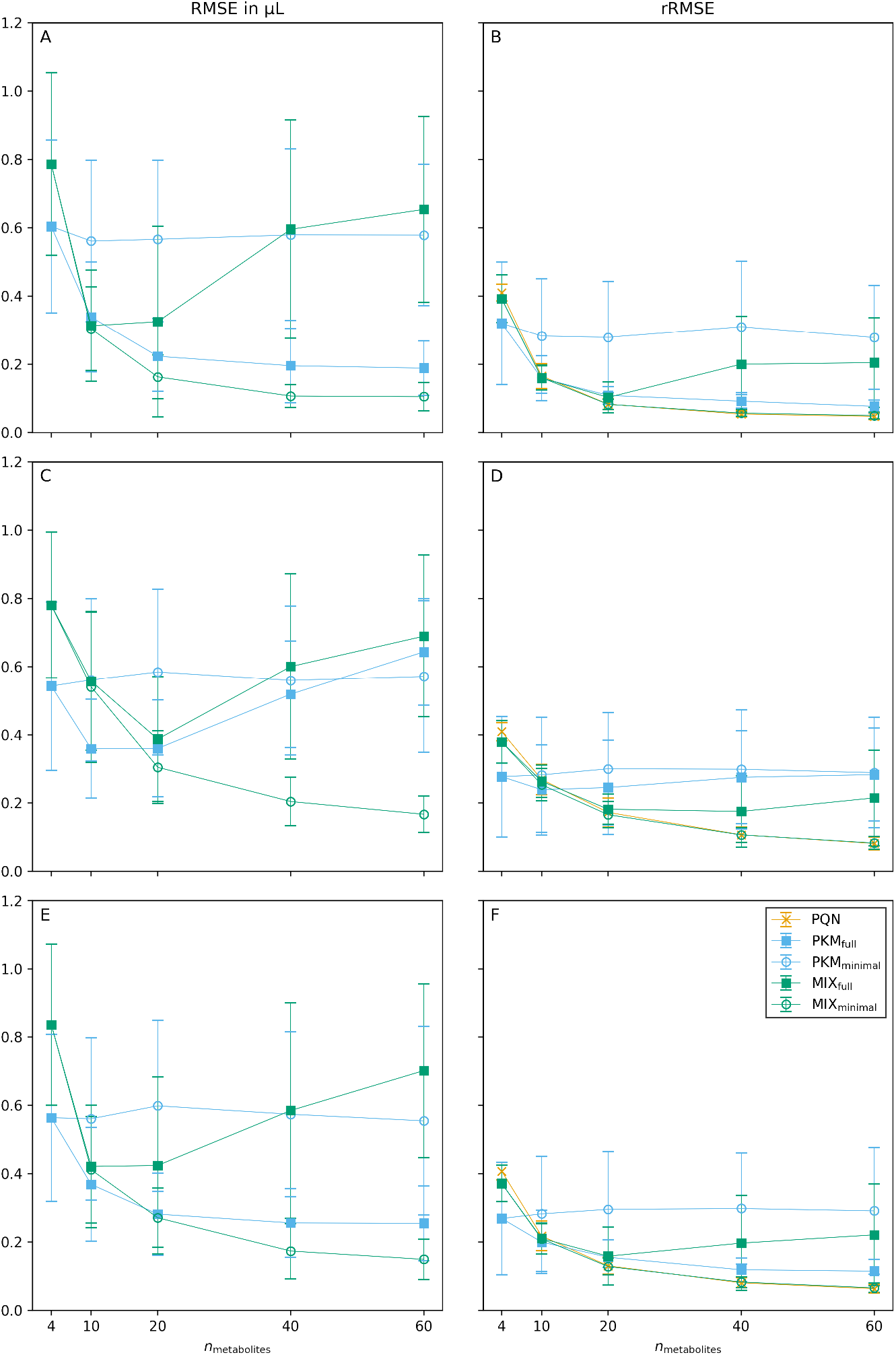
Goodness of normalization measures of synthetic data simulations. The mean for 100 replicates for different sweat volume normalization models is given for RMSE (left column) and rRMSE (right column). Results for simulations v1, v2, and v3 are shown in rows one, two, and three, respectively. The error bars represent standard deviations of the replicates. For the PQN method no RMSE can be calculated.

PKM_full_ which fits a kinetic function through all possible metabolites (ℓ = *n*_metabolites_) performs well (low RMSE, low rRMSE) when the **C** data originates from a kinetic function (simulation v1, Figure 6A, B). However, when the underlying data does not originate from kinetic time series (simulation v2, Figure 6C, D) its performance is reduced drastically. For PKM_full_ this is resembled in an increase of RMSE (from 0.19 ± 0.08 μL to 0.64 ± 0.16 μL for *n*_metabolites_ = 60) as well as of rRMSE (from 0.08 ± 0.02 to 0.28 ± 0.14 for *n*_metabolites_ = 60).

Another observation is the behaviour of PQN. Its rRMSE approaches a value close to 0 with increasing *n*_metabolites_, indifferently on how the underlying data was generated.

Interestingly, the results from simulation v3 lie between the results from simulation v1 and v2. This gets especially evident when comparing the performance of PKMfull in Figure 6. Such a result suggests that not all of the untargeted metabolites measured are completely random, but some can be described with the modified Bateman function. This leads to the hypothesis that after sweat volume normalization, the real finger sweat data (from which values for v3 were sampled) has high potential for the discovery of unknown kinetics.

Exact numbers for RMSE and rRMSE for all normalization methods and *n*_metabolites_ are given in Supplementary Tables S3 and S4 respectively. Moreover, pairwise comparisons of RMSE and rRMSE of normalization methods relative to the results from PKM_minimal_ are plotted in Supplementary Figure S7.

##### Statistical Testing

As at *n*_metabolites_ = 60 the goodness of normalization measures start to flatten out, we further investigated this condition for statistical significance. We used the two-sided non-parameteric Wilcoxon signed-rank test to compare pairwise differences in RMSE and rRMSE between the tested models. p-values for all combinations are given in Supplementary Tables S5 and S6.

As Figure 6 already indicated, the overall best performance in RMSE as well as rRMSE is observed for the MIX_minimal_ model. For *n*_metabolites_ = 60 it significantly outperforms every other method’s RMSE (Figure 7). Moreover, MIX_minimal_’s performance in rRMSE is at least equal to or better than all other tested methods (Supplementary Table S6) with one exception: the comparison of rRMSE of MIX_minimal_ and PQN in simulation v1 shows significant difference (p = 0.0029), however, the absolute values of rRMSE are still very similar (0.049 ± 0.010 and 0.047 ± 0.009 respectively). Compared to the previously used PKM_minimal_ [20], the RMSE of MIX_minimal_ improves by 73 ± 10 %, the rRMSE by 43 ± 12 % (Supplementary Figure S7). Analogously to Figure 7 for simulation v3, the results of simulations v1 and v2 are shown in Supplementary Figure S8 and S9 respectively.

**Figure 7.**
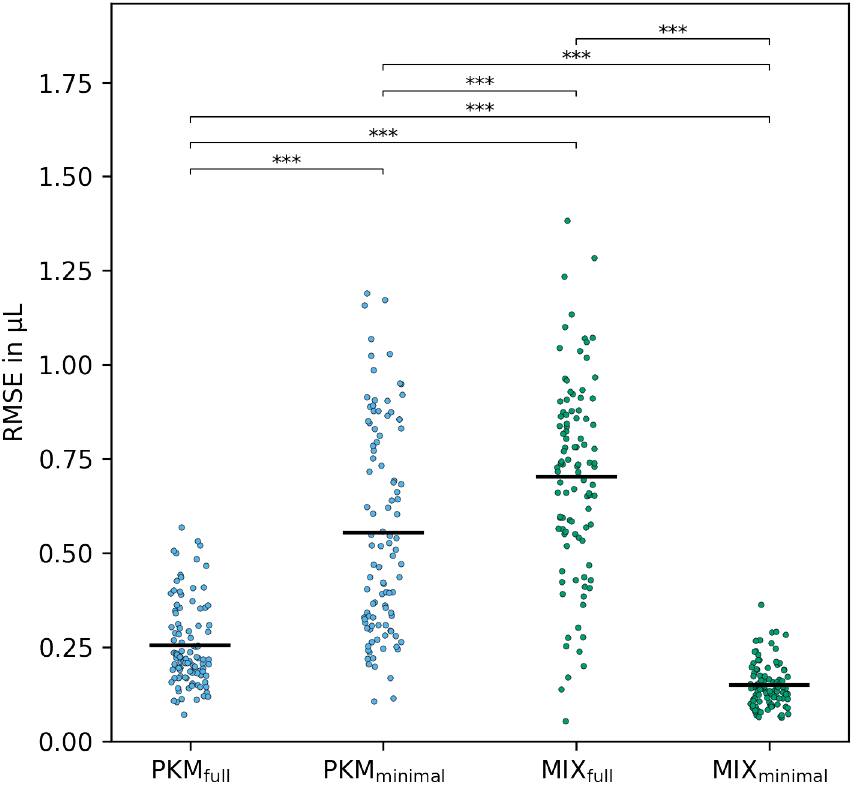
RMSE measures of simulation v3 with *n*_metabolites_ = 60. The significance between the methods was calculated on 100 paired replicates with the two-sided Wilcoxon signed-rank test.

The two-sided version of the Wilcoxon signed-rank test was used to test for any difference in between multiple normalization methods. After it became evident that MIX_minimal_ performed best, we used an one-sided version of the Wilcoxon signed-rank test to verify if RMSE and rRMSE are significantly decreased by MIX_minimal_ compared to all other normalization methods. The resulting p-values are listed in Supplementary Table S7. Again, MIX_minimal_ significantly outperformed all other tested methods in RMSE and rRMSE except for PQN in any of the simulations.

We, therefore, conclude that normalizing the sweat volume by the MIX_minimal_ method reduces the error for the estimated *V* compared to other tested methods. Compared to PKM, MIX_minimal_ has the advantage that its performance does not vary if metabolites’ concentration time series can be described with a modified Bateman function (i.e. simulations v1, v2 v3 have little influence on its performance). Therefore, it is especially advantageous if this property cannot be guaranteed.

#### 4.1.2 Computational Performance

Analysis of metabolomics data sets is usually a computationally exhaustive process. There are several steps in (pre-)processing that need to be executed, many of them lasting for hours. Therefore, computational time can quickly stack to large numbers. Normalization models are no exception to this general rule. As *n*_metabolites_ in a pharmacokinetic model increases, the time for optimization of pharmacokinetic models may become limiting. Therefore, we investigated the average time for one time series normalization for different methods and different numbers of metabolites.

The computational time spent for one optimization step as a function of *n*_metabolites_ is given in Figure 8 for simulation v3. It increased for some normalization models, however not for all of them and not equally. Within the investigated range, PQN stays well under 1 second per normalization, whereas with PKM_full_ the normalization time increases drastically from 1.6 ± 1.1 s for a model with 4 metabolites to 110 ± 44 s for 60 metabolites. Similar normalization times were observed for MIX_full_ maxing out at 19 ± 22 s for *n*_metabolites_ = 60. In stark contrast to the exponential increase in computational power needed for full models are the minimal models. Their time to optimize stays nearly constant (< 3 s) within the investigated metabolite range (Supplementary Table S8).

**Figure 8.**
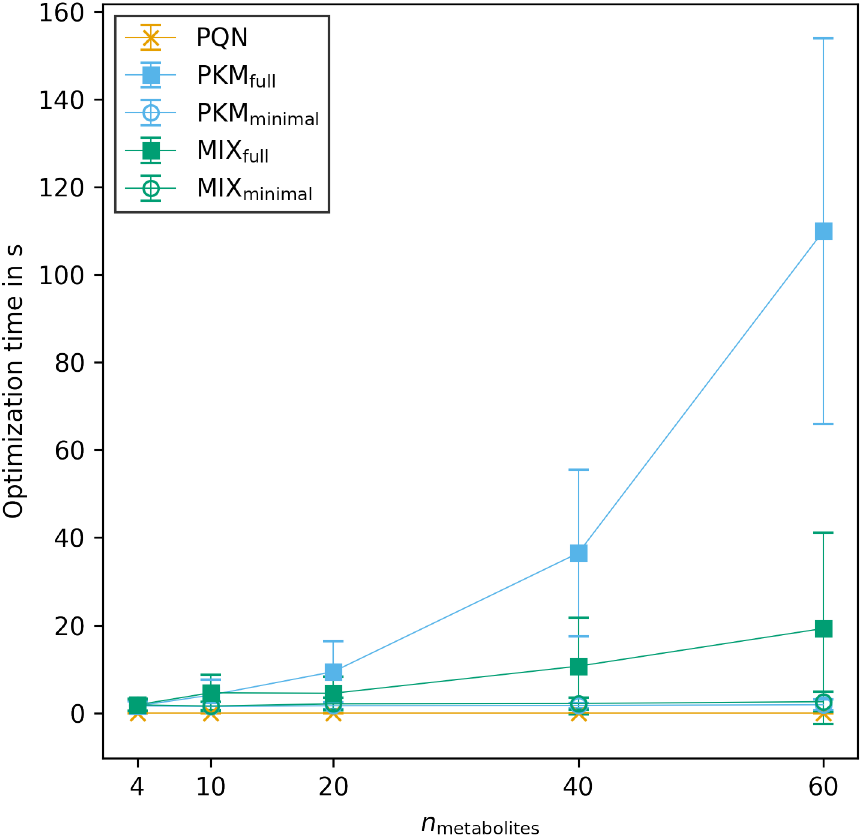
Time in seconds for optimization of one normalization model in simulation v3. The error bars represent the standard deviation of normalization times between 100 replicates.

Here we demonstrate that MIX_minimal_ is not only superior to other tested models in terms of its normalization performance, but also in terms of computational feasibility. We hypothesize that even data sets with thousands of untargeted metabolites will have a minor impact on its speed.

### 4.2 Comparison of PQN and MIX

#### 4.2.1 Influence of Noise on PQN

In untargeted metabolomics it is often difficult to distinguish between metabolites originating from the actual matrix of interest or from contamination. As PQN includes all untargeted metabolites in its calculation, metabolites stemming from contamination might become a problem as their fold change is independent on the sweat volume, which changes the underlying distributions of quotients. Therefore, we investigated the influence of different fractions of metabolites originating from contamination (i.e. noisy data). Furthermore, we tested if scaling of Q^PQN^ values can counteract errors introduced by noise.

Figure 9A demonstrates the problem of using the probabilistic quotient normalization on noisy raw data. The direction of size effects can still be explained when noise is present, however, absolute values of the size effects decrease. Thus, in Figure 9A the coefficient of variation (i.e. the standard deviation over the mean) of **Q**^PQN^ is a measure for the average value of the estimated size effect over one synthetically generated time series. As the fraction of noise (*f_n_*, X-axis) increases the coefficient of variation decreases drastically and approaches 0 when *f_n_* → 1.

**Figure 9.**
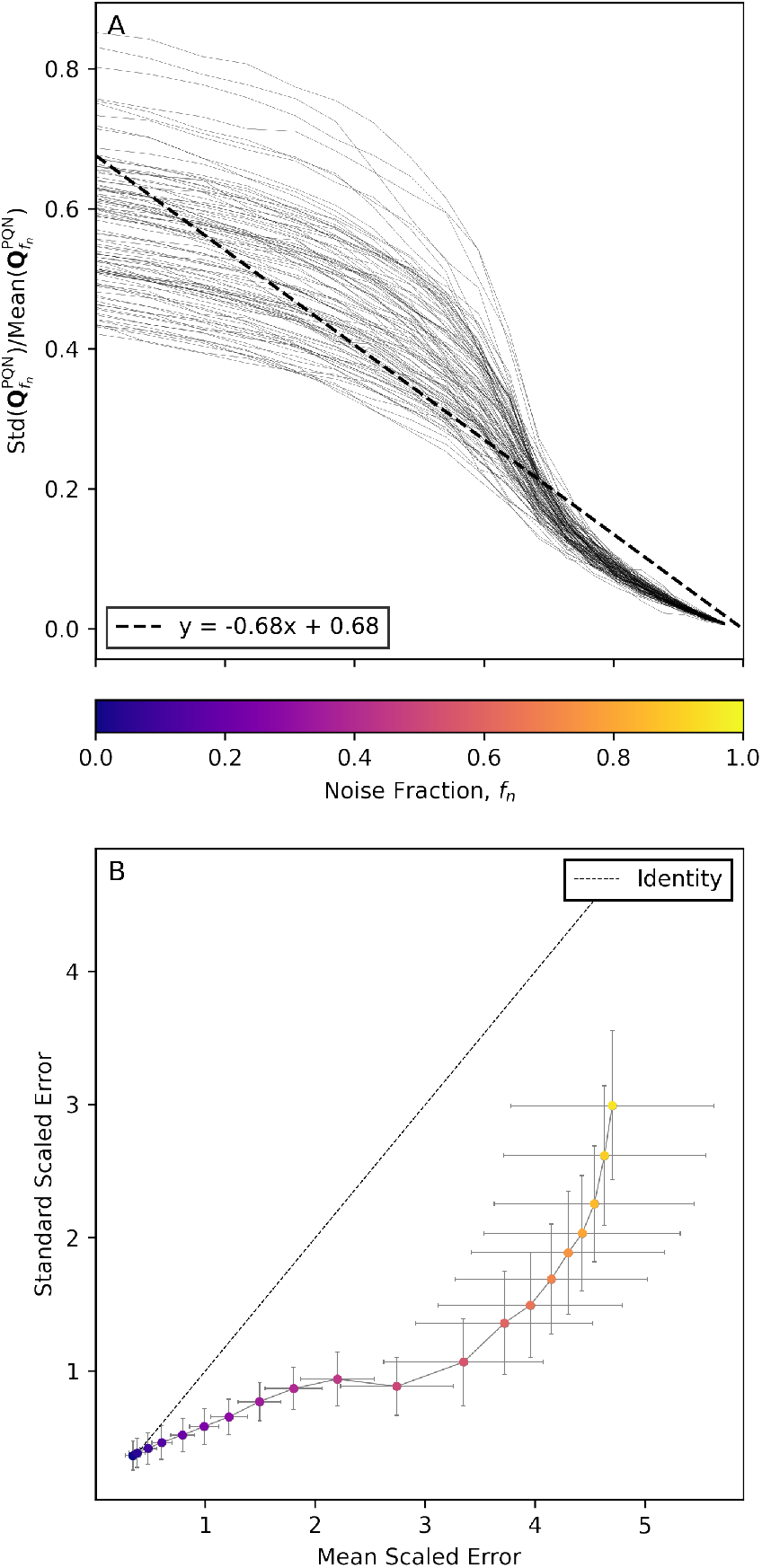
Influence of the fraction of noisy data on the error of PQN calculation. Panel A illustrates the change of the coefficient of variance of Q^PQN^ (Y-axis) as the noise fraction (*f_n_*, Y-axis with the same tick labels as the color bar) increases. Panel B shows the error size of calculated Q^PQN^ to true **V** with mean scaling (X-axis) and standard scaling (Y-axis). The color of points relates to the noise fraction as depicted in the color bar.

Figure 9B shows the performance of scaling methods to counteract the reduction of coefficient of variation as described above. The mean scaled error (X-axis) and standard scaled error (Y-axis) as calculated by Equations 17 are plotted against each other. When *f_n_* ≤ 0.05, mean scaling outperforms standard scaling, however, thereafter the standard scaled **Q**^PQN^ is less erroneous than the mean scaled version.

When incorporating **Q**^PQN^ values to the MIX model it is important to correct for errors introduced by noise. As this result shows that standard scaling reduces the detrimental effect of noise on the calculation of **Q**^PQN^, we used standard scaling throughout the study for MIX normalization. Moreover, this result underlines the good performance of standard scaling in biological data sets [43].

#### 4.2.2 Synthetic Data Simulations with Noise

The synthetic data used for the analysis of Section 4.1 did not contain any metabolites that are classified as noise, i.e. their 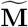 is not influenced by size effects (Equation 15). This, however, is not necessarily a realistic assumption as there are many sources of contaminants in metabolome measurements. Noisy metabolites can be either introduced by biological means (e.g. metabolites that do not originate from sweat but from the surface of the skin in sweat measurements) [44] or by experimental handling [45]. As shown in Figure 9, this noise in data negatively affects the performance of PQN. Thus, the goodness of PQN in the results of Section 4.1 is probably overestimated.

To get a more accurate view on the goodness of normalization of PQN and MIX_minimal_, we tested their performance on synthetic data with different fractions of noise, *f_n_.* In order to do so, we created 100 replicates of synthetic data sampled from real data (i.e. simulation v3) for 10 equidistant noise fractions ranging from *f_n_* = 0 to *f_n_* = 0.9 with *n*_metabolites_ = 60. In all simulated data, only untargeted metabolites were affected by the introduction of noise, as we assumed that for targeted metabolites (i.e. ℓ = 4) with known pharmacokinetic behaviour one can be highly confident that the measurements are not originating from contaminants.

The rRMSE of PQN and MIX_minimal_ is plotted in Figure 10. Only when zero noise was present in the synthetic data set, MIX_minimal_ did not improve upon PQN, however as the fraction of noise increased, MIX_minimal_ significantly outperformed PQN in terms of rRMSE. The *p*-values for all noise fractions are listed in the Supplementary Table S9.

**Figure 10.**
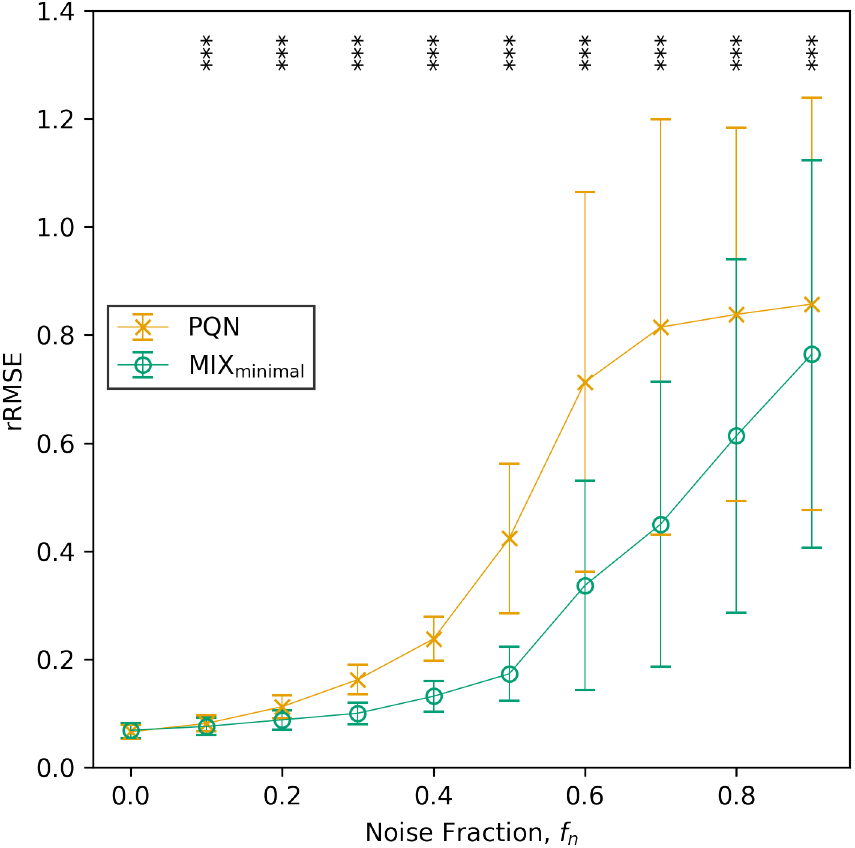
Comparison of the rRMSE of PQN and MIX_minimal_ on data with different fractions of noise. Significant differences in rRMSE between PQN and MIX_minimal_ were tested with an one-sided pairwise Wilcoxon signed-rank test.

The difference of rRMSE between PQN and MIX_minimal_ in Figure 10 is related to the difference of mean and standard scaled errors in Figure 9B. PQN alone cannot utilize the improved performance of standard scaling as Std(T(**V**)) has to be known for its calculation (Equation 17b). However, when normalizing with MIX_minimal_, Std(*T*(**V**)) can be estimated from the pharmacokinetic part of the model (Equation 9c) significantly improving its quality.

### 4.3 Application to Real Data

#### 4.3.1 Caffeine Network

Previously, we identified and quantified four metabolites (caffeine, paraxanthine, theobromine, and theophylline) in a time series after the ingestion of a single dose of caffeine [20]. To investigate the performance of normalization models on a real finger sweat data set, we split all measured 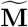 time series into three parts that contained pairs of targeted metabolites each, only one shared by all, namely caffeine (compare Figure 4 top and bottom network). Subsequently we fitted a PKM_minimal_ and MIX_minimal_ models (ℓ = 2) with adapted kinetics (Methods Section 3.4) through the three sub data sets. Due to the nature of the metabolite subnetworks (Figure 4 bottom panel) it is possible to calculate two kinetic constants describing the absorption and elimination of caffeine (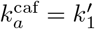 and 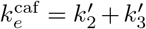) in all three cases. As the data for all three subnetworks was measured in the same experiment we can assume that the underlying ground truth of these constants has to be the same. Therefore, by comparing the standard deviation of kinetic constants it is possible to infer the performance of normalization methods.

In panels A and B of Figure 11 the standard deviations of fitted kinetic constants within one measured 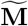 time series are illustrated. Panel A shows that the standard deviations of the absorption constant of caffeine, 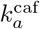, of PKM_minimal_ are significantly larger than of the MIX_minimal_ model (*p* = 5.8 × 10^-4^, *n* = 37, one-sided Wilcoxon signed-rank test). Likewise, a significant decrease of the size of standard deviations of MIX_minimal_ was found compared to the previously published PKM_minimal_ model (*p* = 1.5 10^-5^) for the constant of caffeine elimination, 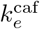 (panel B, Figure 11).

**Figure 11.**
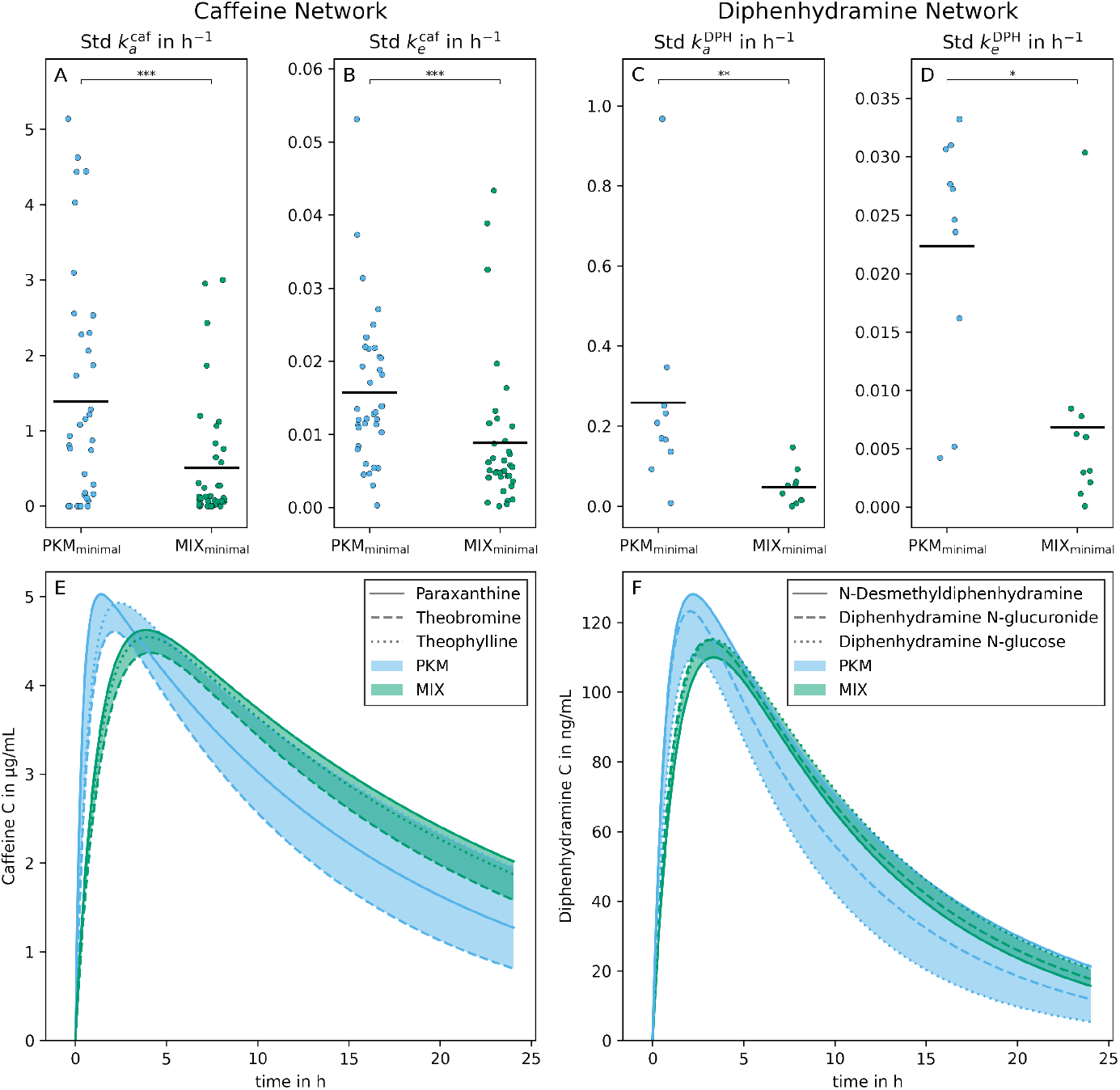
Method validation with finger sweat (left column) and blood plasma (right column) data from Brunmair et al., 2021 [20] and Panitchpakdi et al., 2021 [40] respectively. On panels A to D, the standard deviations of constants of absorption and elimination of caffeine and diphenhydramine 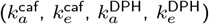 between the three modeled subnetworks are plotted. The number of points per method corresponds to the number of concentrations time series present in both data sets (i.e. 37 and 10 for sweat and plasma respectively). A one-sided Wilcoxon signed-rank test was used to test for significant differences. Panels E and F show the estimated concentration time series of caffeine and DPH plotted from the three different subnetworks. The lines are named after the second metabolite with known kinetic present in the subnetwork, however they all refer to **C** of caffeine and DPH themselves. The colors of curves and the area between them indicate the results from normalization with PKM_minimal_ or MIX_minimal_ respectively.

In panel E of Figure 11 one exemplified normalized C time series of caffeine in sweat is depicted as fitted for all three subnetworks with PKM_minimal_ and MIX_minimal_ respectively. The selected time series illustrates the median of differences in standard deviations between PKM_minimal_ and MIX_minimal_ from panels A and B of Figure 11. The area enclosed by the *C*s of MIX_minimal_ models is smaller than from PKM_minimal_.

We emphasize that in our original study the caffeine degradation directly produces paraxanthine, theobromine, and theophylline, thus pharmacokinetic parameters *k*_2_, *k*_3_, *k*_4_ are explicitly linked [20]. Therefore, the kinetic network resembled specific kinetics of that metabolic pathway (Figure 4 top panel). In contrast, in previous sections we assumed that the underlying pathway structure is not known. Thus parameters are not linked, which implies that parameters are less constrained. Yet in this section, we demonstrated that the fundamental improvement found by switching from PKM to a MIX model can be also translated back again to a more specific metabolic network (Figure 4 bottom panel). In order to support this argument, we show the applicability of the MIX_minimal_ normalization method on a real finger sweat data set. The results with real data emphasize the validity of the simulations done on synthetic data sets. They show that, especially when known metabolic networks are small, the MIX_minimal_ model significantly improves the robustness of normalization and thus kinetic constants inferred from finger sweat time series measurements.

#### 4.3.2 Diphenhydramine Network

In the original study [40] the authors measured time series abundances in the blood plasma after the application of a single dose of diphenhydramine (DPH). 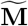 from targeted DPH (known pharmacological constants, known kinetics) as well as untargeted metabolization products (N-desmethyl-DPH, DPH N-glucuronide, DPH-glucose, known kinetics) and several other untargeted metabolites (unknown kinetics) were reported. Similar to sweat, although less pronounced, plasma also suffers from size effects (i.e. a systematic error in the measurements) introduced by biological means or preanalytical sample handling [46, 47]. Thus, we used the reported data as a second real data set for validation of the performance of MIX_minimal_. The validation was performed in analogy to the caffeine study where a full network (Supplementary Figure S4) is split into three subnetworks (Supplementary Figure S5, for details see Methods Section 3.5).

In panels C and D of Figure 11 the standard deviations of fitted kinetic constants within one measured 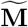 and three fitted subnetworks are illustrated. Again, the standard deviations of 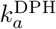 of PKM_minimal_ are significantly larger than of MIX_minimal_ (*p* = 2,0 × 10^-3^, *n* = 10, one-sided Wilcoxon signed-rank test, panel C). A similar significant decrease of the standard deviations are also found for 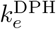 (*p* = 3.2 10^-2^, panel D).

In panel F of Figure 11 one exemplified, normalized *C* time series of DPH in plasma is depicted as fitted for all three subnetworks with PKM_minimal_ and MIX_minimal_ respectively. The time series was selected as it is closest to the median of the differences in standard deviations between PKM_minimal_ and MIX_minimal_. It is visible that the area enclosed by the *C* resulting from the MIX_minimal_ model is smaller than from PKM_minimal_.

This validation illustrates the performance of the normalization models presented in this study on a data set that was measured independently from the development of said methods. The results of the plasma validation study are similar to the results observed for the finger sweat study; again, MIX_minimal_ improves the robustness (i.e. reduces standard deviations) of size effect normalization.

Even though there is a significant decrease in the standard deviation of 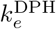 with MIX_minimal_ compared to PKM_minimal_, MIX_minimal_ also produced an outlier (Figure 11D). The reason for this outlier is that on rare occasions MIX_minimal_ is not able to detect any size effects due to convergence issues (Supplementary Figure S10A). To investigate this results we performed synthetic data simulations (Supplementary Figure S10B). There, we found that this behaviour of MIX_minimal_ can be observed when two different **V** vectors are applied to ℓ and ℓ+ metabolites. Therefore, we hypothesize that the clearly visible malfunction of MIX_minimal_ to detect size effects (i.e. the variance of estimated **V** is close to 0) gives an indication to scientists that size effects might not be a major concern in such a data set. In this specific blood plasma time series measurement, for example, the size effects might have been too small compared to other error sources to be identified by MIX_minimal_.

To summarize, with this validation we show that the generalized normalization models, as implemented in this study can directly be used for the normalization of real data as long as the modified Bateman function is able to describe the measured kinetics reasonably well and size effects are large enough to be detectable.

## 5 Discussion

In this study we present a generalized framework for the PKM normalization model, first introduced in reference [20]. Moreover, we extend the existing model to incorporate untargeted metabolite information, dubbed as MIX model. Both models are implemented in Python and are available at GitHub https://github.com/Gotsmy/sweat_normalization.

The quality of normalization methods was tested on synthetic data sets. Synthetic data sets are necessary as it is impossible to obtain validation data without fundamentally changing the (finger) sweat sampling method as described above [20]. However, three different synthetic data generation methods (v1, v2, v3) were employed to ensure that synthetic data sets are as close to real data as possible. We found that, when *n*_metabolites_ ≥ 60, MIX_minimal_ performs equally well or better than all other tested normalization methods.

Despite true *V* values remaining unknown, the real finger sweat data can be used as validation for relative robustness of normalization methods. There, MIX_minimal_ significantly outperforms PKM_minimal_. The decreased variance of kinetic constants estimated by MIX_minimal_ likely originates from the fact that **Q**^PQN^ does not differ much for three subsets as long as sufficiently many *n*_metabolites_ = 60 are present in each subset. On the other hand, as only few data points are used for PKM_minimal_ optimization, small errors in one of the two targeted metabolites measured mass have a high potential to change the normalization result.

Additionally, the performance of PKM_minimal_ and MIX_minimal_ were compared on a blood plasma data set taken from a study independent from any measurements used for the development of the normalization models. There, we were able to demonstrate the same improvement from PKM_minimal_ to MIX_minimal_ in normalization robustness. Moreover, we show that the generalized normalization models as implemented as Python class in this study can be easily used for size effect normalization with little additional coding necessary.

To recapitulate, the proposed MIX_minimal_ model has several crucial advantages over other tested methods.
- MIX_minimal_ significantly outperforms PKM_minimal_ in relative (rRMSE, −43 ± 12 %) and absolute (RMSE, −73 ± 10 %) errors with as little as 60 untargeted metabolites used as additional information (Figure 7).
- MIX_minimal_ is invariant to whether untargeted metabolites follow an easily describable kinetic concentration curve (Figure 6).
- Without noise, MIX_minimal_ performs equally well as PQN for relative abundances, but additionally it estimates absolute values of *V*, similar to pharmacokinetic (PKM) models (Figure 6).
- When noise is present MIX_minimal_ also outperforms PQN for relative abundances (Figure 10).
- MIX_minimal_ performs well in this proof of principle study, moreover, it may be used as a basis for further improvements. Firstly, different, more sophisticated statistical normalization methods (e.g. EigenMS [27]) could be used as input for the PQN part of the model. Secondly, Bayesian priors describing uncertainties of different metabolites could be implemented over the λ parameter in a similar fashion as discussed in reference [48].
- Strikingly, the results showed that for all normalization methods tested the RMSE and rRMSE values flattened once 60 metabolites were present in the original information. This suggested that the presented normalization models, especially MIX_minimal_ can be applied even for biomatrices or analytical methods with as few as 60 compounds measured.
- Although MIX_minimal_ was developed especially with sweat volume normalization in mind, it can easily adapted for other biomatrices, e.g. plasma (Figure 11).

## 6 Conclusion

In this study we described and defined the MIX metabolomics time series normalization model and compared it to PKM. Subsequently, we elaborated several advantages of the MIX_minimal_ model over PKM and previously published normalization methods. We are confident that this will further improve the reliability of metabolomic studies done on finger sweat and other conventional and non-conventional biofluids. However, we acknowledge that a more thorough investigation with data sets of several more quantified metabolites and determined sweat volumes need to be carried out to assess the full potential of the proposed method.

## Declarations

Ethics approval and consent to participate

Not applicable.

## Consent for publication

Not applicable.

## Availability of data and materials

All analysis (except stated otherwise) was performed in Python 3.7 heavily relying on NumPy [49], Pandas [50], and SciPy [32]. Code for simulations, scripts for creation of figures and original and generated data is available on GitHub https://github.com/Gotsmy/sweat_normalization under the GNU GPL version 3 license.

## Authors’ contributions

Conceptualization was done by MG and JZ. Funding was aquired by CG and JZ. Methodology was developed by MG and JB. Software was developed by MG. Formal analysis was done by MG and CB. Investigation was done by MG and JB. Resources were allocated by CG and JZ. Validation was done by MG, JB, and CB. The study was supervised by CG and JZ. Visualization was done by MG. Original draft was written by MG and JZ. Review and editing of the manuscript was done by all authors.

## Acknowledgements

We thank Morgan Panitchpakdi and Shirley Tsunoda for their help in interpreting the plasma data set [40].

## Authors’ information

Not applicable.

## Abbreviations

| Symbol | Name |
| --- | --- |
| $a_{\text{sample}}$ | sampling skin area |
| $b$ | part of modified Bateman function |
| $C, \mathbf{C}$ | underlying concentration (vector) |
| $c_0$ | kinetic parameter |
| $d$ | kinetic parameter |
| $F$ | modified Bateman function |
| $f_n$ | noise fraction |
| $V^{\text{ref}}$ | PQN correction factor |
| $i$ | time point index |
| $j$ | metabolite index |
| $k$ | kinetic parameter |
| $\ell$ | metabolites used for kinetic fitting |
| $\ell+$ | metabolites not used for kinetic fitting |
| $\mathcal{L}$ | loss |
| $L$ | loss function |
| $lag$ | kinetic parameter |
| $\widetilde{M}, \widetilde{\mathbf{M}}$ | measured mass (vector) |
| $M^{\text{ref}}$ | reference mass for PQN |
| $m/z$ | mass over charge ratio |
| $n_{\text{metabolites}}$ | number of metabolites |
| $n_{\text{time points}}$ | number of time points |
| $p$ | $p$-value |
| $q_{\text{sweat}}$ | sweat rate |
| $Q^C$ | median concentration fold change of two samples |
| $Q^M$ | median mass fold change of two samples |
| $Q^{\text{PQN}}, \mathbf{Q}^{\text{PQN}}$ | normalization quotient (vector) calculated by PQN |
| $R^2$ | coefficient of determination |
| rRMSE | relative measure of goodness of normalization |
| RMSE | absolute measure of goodness of normalization |
| Std | standard deviation |
| $T$ | transformation function |
| $t$ | time |
| $V, \mathbf{V}$ | collected (sweat) volume (vector) |
| Var | variance |
| v1, v2, v3 | synthetic data sets |
| $Z$ | scaling function |
| $\epsilon$ | experimental error vector |
| $\theta$ | kinetic parameter vector for fitting |
| $\lambda$ | loss weighting parameter |
| $\tau$ | time to collect one sample |

