## Supplementary Information for "Probabilistic quotient’s work & pharmacokinetics’ contribution: countering size effect in metabolic time series measurements"

#### 1 Supplementary Figures

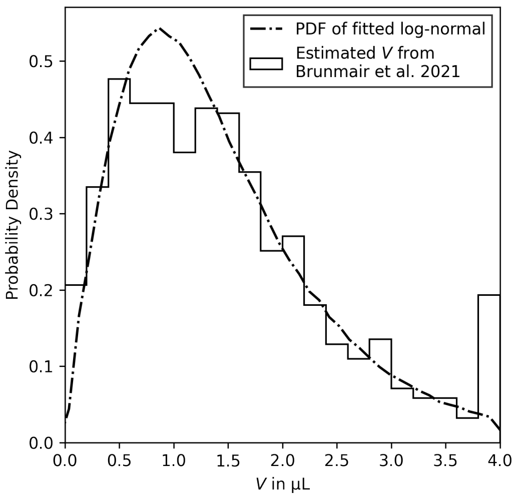


**Figure S1 Probability distribution of $V$.** Distribution of $V$ values estimated in ref. [20] (histogram) and sampled log-normal distribution for synthetic data generation (dashed line).

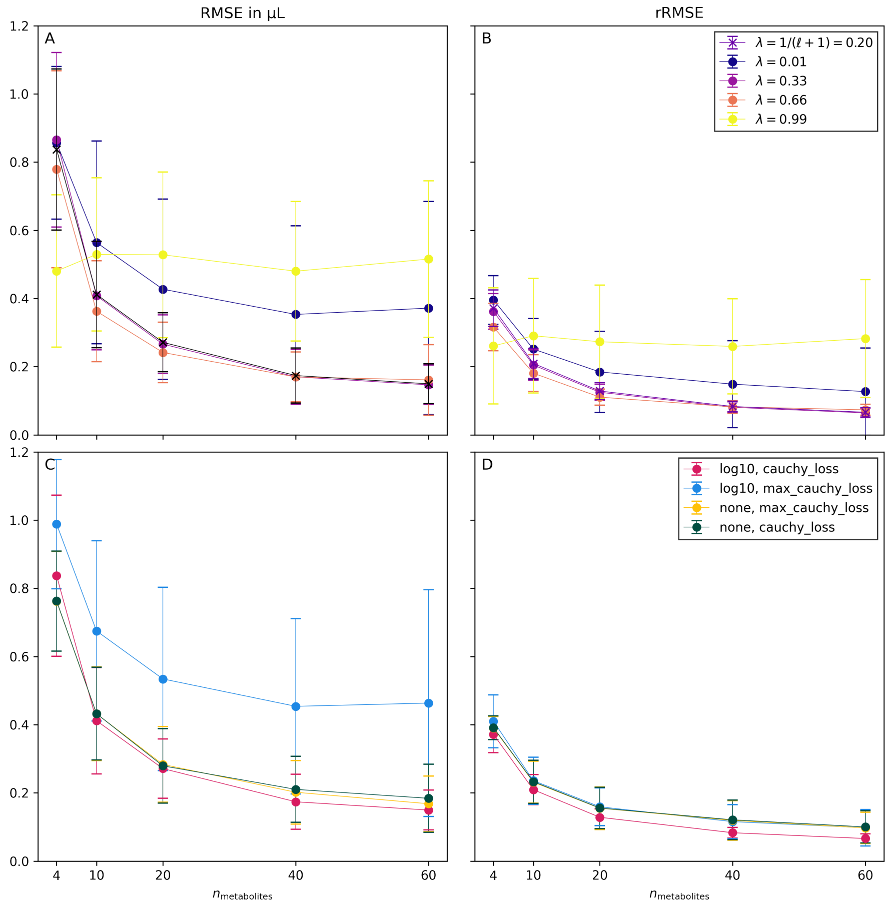


**Figure S2 Testing the influence of hyperparameters on the performance of MIX$_{\text{minimal}}$** Simulations were performed for 100 replicates with $n_{\text{metabolites}} = 60$ and synthetic data from simulation v3. Panels A and B show the RMSE and rRMSE of MIX$_{\text{minimal}}$ with different values for the weighting parameter $\lambda$. The goodness of normalization by MIX$_{\text{minimal}}$ is insensitive to a wide range of values for $\lambda$. Panels C and D show the RMSE and rRMSE of different combinations of transformation functions $(T)$ and loss functions $(L)$. It is visible that the combination of `log10` transformation function and `cauchy_loss` loss function result in the lowest RMSE and rRMSE in the MIX$_{\text{minimal}}$ model. Thus they were selected for all further analysis.

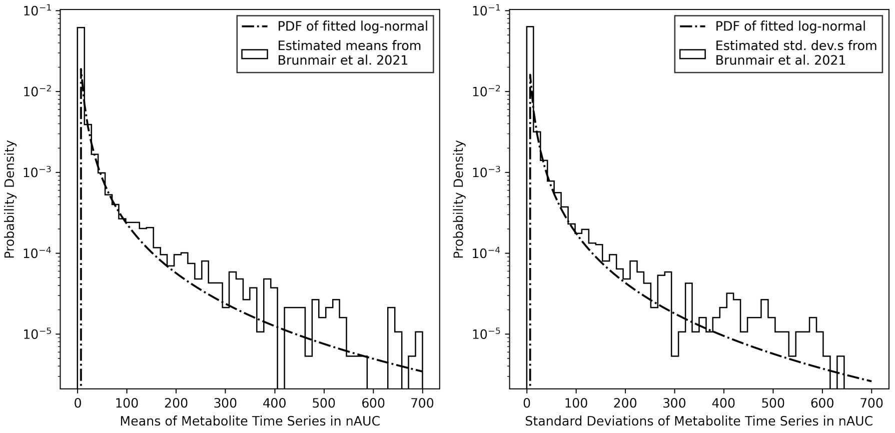


**Figure S3 Probability distribution of measured abundances.** Distribution of means (left panel) and standard deviations (right panel) of metabolites per time series from real finger sweat data [20] (histogram) and sampled distribution for synthetic data generation (dashed line).

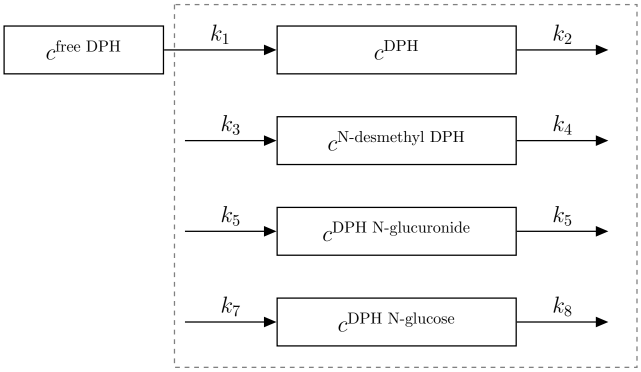


**Figure S4 Metabolic network of diphenhydramine (DPH), N-desmethyl-DPH, DPH N-glucuronide, and DPH N-glucose absorption and elimination.** The system boundary (dashed line) represents the human body.

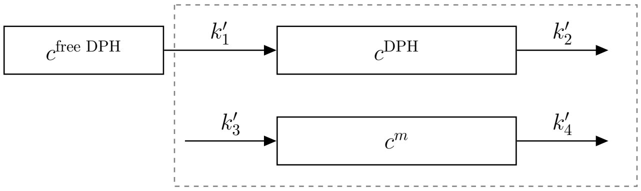


**Figure S5 Simplified metabolic network of diphenhydramine (DPH) and $m$ absorption and elimination** ($m \in$ {N-desmethyl-DPH, DPH N-glucuronide, and DPH N-glucose}). The system boundary (dashed line) represents the human body.

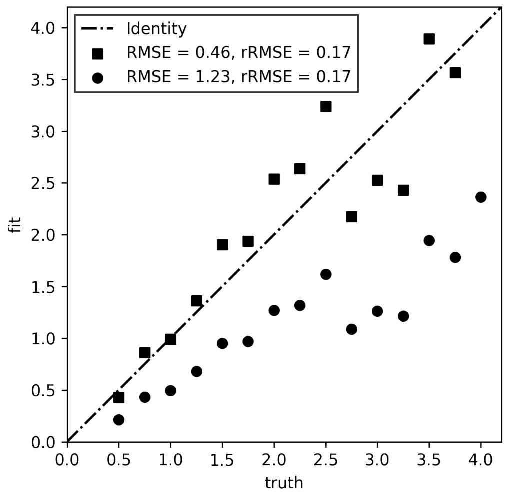


**Figure S6 Example of RMSE and rRMSE applied to two fits (squares and circles).** The error multiplied to both fits is exactly the same, however, the circles are additionally scaled by a factor of 0.5. As their relative size is not changed by a scaling the rRMSE is also unaffected. However, the scaling does have an influence on the absolute offset to the truth, thus an increased RMSE.

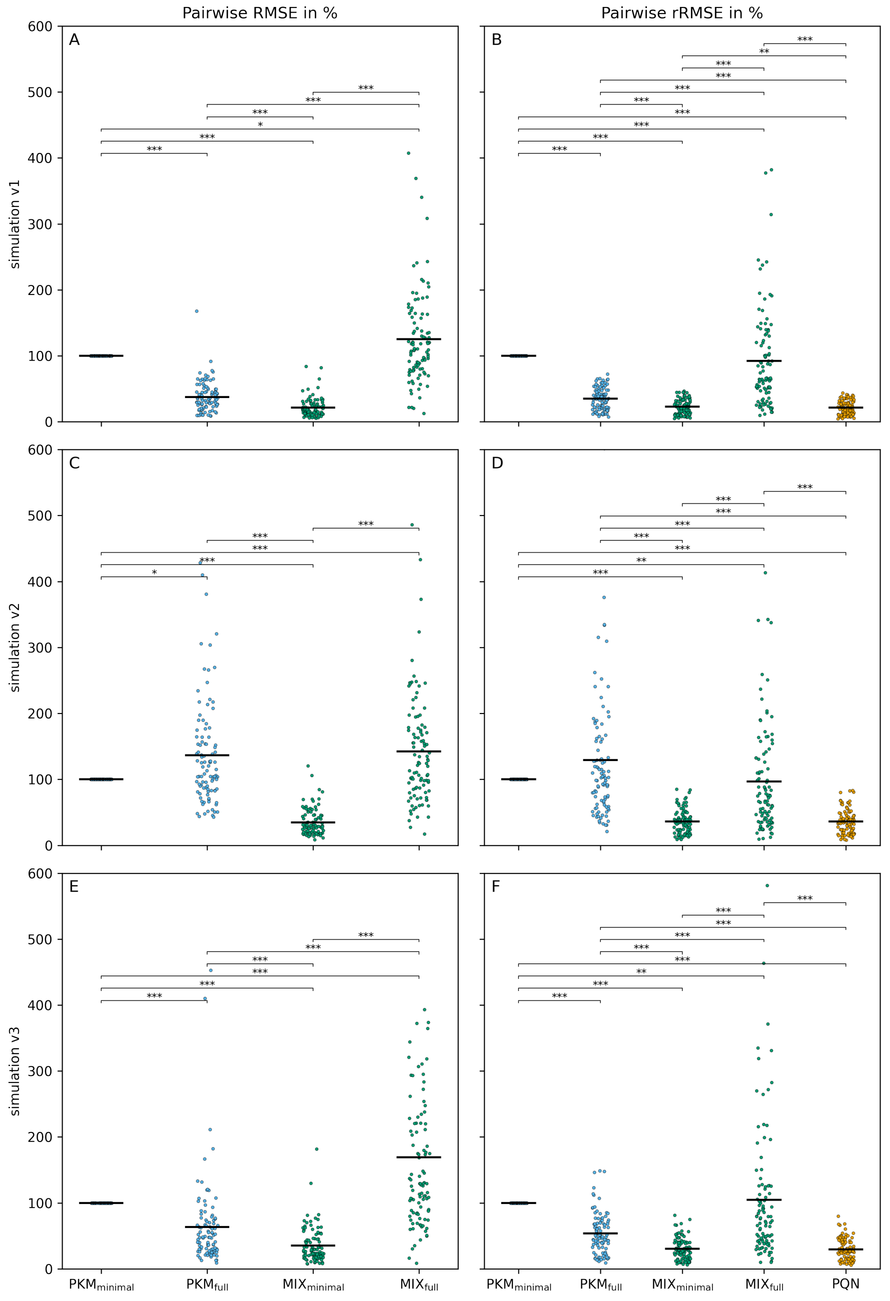


**Figure S7 Pairwise comparison of the RMSE (left column) and rRMSE (right column)** of 100 replicates of synthetic data sets from simulations v1 (first row), v2 (second row), and v3 (third row) normalized to the results from $PKM_{minimal}$. The significance between the methods was calculated on the absolute RMSE/rRMSE of 100 paired replicates with the two-sided Wilcoxon signed-rank test.

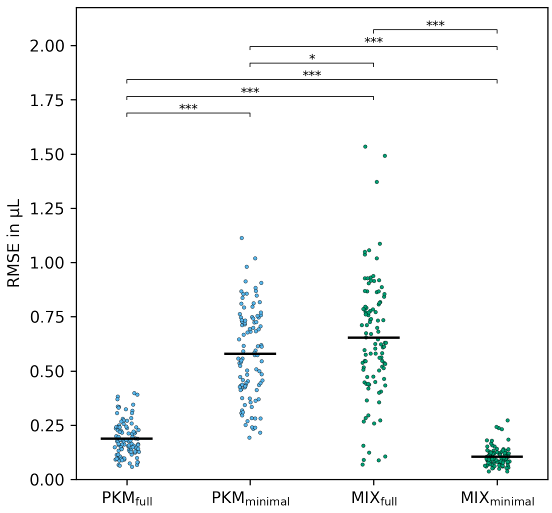


**Figure S8 Results of simulation v1 with $n_{\text{metabolites}} = 60$.** The significance between the methods was calculated on 100 paired replicates with the Wilcoxon singed-rank test.

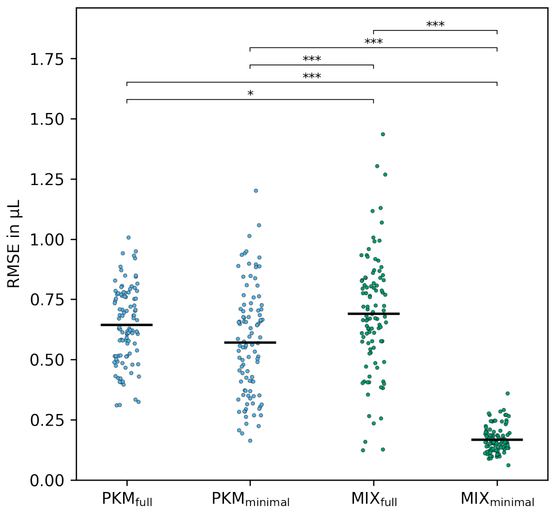


**Figure S9 Results of simulation v2 with $n_{\text{metabolites}} = 60$.** The significance between the methods was calculated on 100 paired replicates with the Wilcoxon signed-rank test.

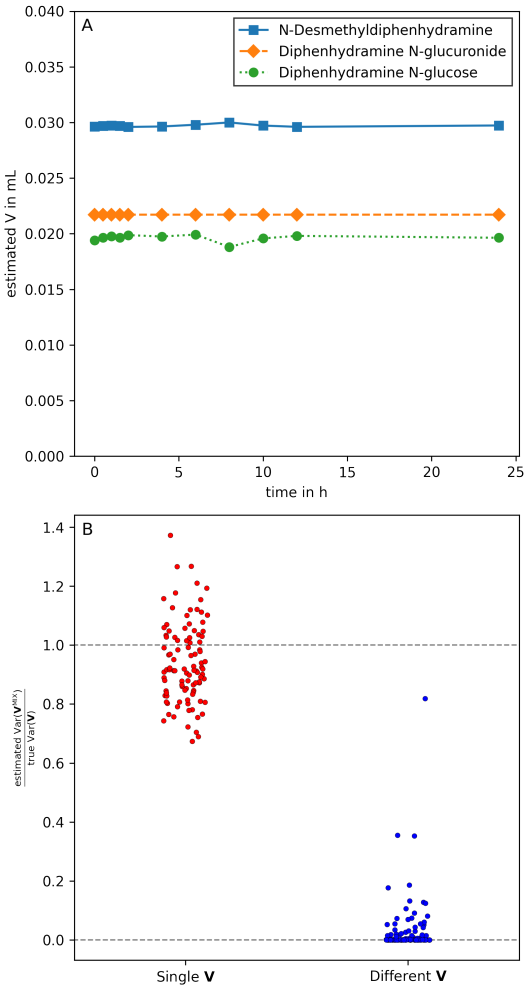


**Figure S10 Example of the estimated $\mathbf{V}$ for the plasma time series of subject 191009.** Panel A shows that instead of discerning size effects, all volumes of the time series converge to one average value for all three subnetworks each (the legends refer to the second metabolite with known pharmacokinetics present in the subnetwork). Panel B shows the variance of $\mathbf{V}$ as estimated by $\text{MIX}_{\text{minimal}}$ divided by the true variance of $\mathbf{V}$ used for synthetic data generation. The synthetic data was generated for 100 replicates, with $n_{\text{metabolites}} = 100$ and simulation v3. For results on the left hand side (red points), a single array of $\mathbf{V}$ was sampled and applied to all metabolites of the data set. In contrast, for results on the right hand side (blue points), two different $\mathbf{V}$ arrays were generated and applied to targeted ($\ell$) and untargeted ($\ell+$) metabolites separately. This was done to test the behaviour of the $\text{MIX}_{\text{minimal}}$ model if the underlying assumption that all metabolites are influenced by the same size effects is violated. The variance of the control simulations (red) over the true variance of data is close to 1, which means that estimated and true variance are similar in size. However, if the assumption of common size effects is violated, the variance of estimated $\mathbf{V}$ by $\text{MIX}_{\text{minimal}}$ approached 0 (blue points). The exact same observation can be made for the results of subject 191009, panel A.

#### 2 Supplementary Tables

**Table S1 Properties of PKM and MIX models used in this study.** The difference in the number of data points originates from the additional $\mathbf{Q}^{\text{PQN}}$ vector used for optimization of MIX.

| model name | PKM | MIX |
|---|---|---|
| model Python class | `PKM_model` | `MIX_model` |
| number of parameters | $n_{\text{metabolites}} \cdot 5 + n_{\text{time points}}$ | $n_{\text{metabolites}} \cdot 5 + n_{\text{time points}}$ |
| number of data points | $n_{\text{metabolites}} \cdot n_{\text{time points}}$ | $(n_{\text{metabolites}} + 1) \cdot n_{\text{time points}}$ |
| loss function, $L$ | `max_cauchy_loss` | `cauchy_loss` |
| transformation function, $T$ | `none` | `log10` |
| scaling function, $Z$ | - | `standard` |
| weighting, $\lambda$ | - | $1/(\ell + 1)$ |

**Table S2 Kinetic parameters used for the toy model.**

| Metabolite | $k_a$ | $k_e$ | $c_0$ | $lag$ | $d$ |
|---|---|---|---|---|---|
| #1 | 2 | 0.1 | 1 | 0 | 0.1 |
| #2 | 2 | 0.1 | 2 | 0 | 0.1 |
| #3 | 2 | 0.1 | 3 | 0 | 0.1 |
| #4 | 2 | 0.1 | 0.5 | 0 | 0.1 |

**Table S3 Mean and standard deviation of RMSE results of all simulations and $n_{\text{metabolites}}$.**

| | Sim. | Mean | Std | Mean | Std | Mean | Std | Mean | Std | Mean | Std |
|---|---|---|---|---|---|---|---|---|---|---|---|
| $n_{\text{metabolites}}$ | - | 4 | | 10 | | 20 | | 40 | | 60 | |
| $\text{PKM}_{\text{full}}$ | v1 | **0.604** | 0.253 | 0.338 | 0.161 | 0.223 | 0.102 | 0.195 | 0.109 | 0.188 | 0.081 |
| $\text{PKM}_{\text{minimal}}$ | v1 | **0.604** | 0.253 | 0.561 | 0.237 | 0.566 | 0.231 | 0.579 | 0.251 | 0.579 | 0.207 |
| $\text{MIX}_{\text{full}}$ | v1 | 0.786 | 0.268 | 0.313 | 0.163 | 0.325 | 0.280 | 0.596 | 0.319 | 0.654 | 0.273 |
| $\text{MIX}_{\text{minimal}}$ | v1 | 0.786 | 0.268 | **0.303** | 0.123 | **0.163** | 0.064 | **0.107** | 0.033 | **0.105** | 0.042 |
| $\text{PKM}_{\text{full}}$ | v2 | **0.543** | 0.247 | **0.359** | 0.145 | 0.360 | 0.142 | 0.519 | 0.157 | 0.643 | 0.156 |
| $\text{PKM}_{\text{minimal}}$ | v2 | **0.543** | 0.247 | 0.561 | 0.237 | 0.584 | 0.243 | 0.560 | 0.218 | 0.571 | 0.221 |
| $\text{MIX}_{\text{full}}$ | v2 | 0.780 | 0.213 | 0.557 | 0.202 | 0.387 | 0.183 | 0.600 | 0.272 | 0.689 | 0.237 |
| $\text{MIX}_{\text{minimal}}$ | v2 | 0.780 | 0.213 | 0.540 | 0.222 | **0.305** | 0.106 | **0.205** | 0.071 | **0.167** | 0.054 |
| $\text{PKM}_{\text{full}}$ | v3 | **0.564** | 0.244 | **0.369** | 0.166 | 0.282 | 0.120 | 0.256 | 0.100 | 0.255 | 0.110 |
| $\text{PKM}_{\text{minimal}}$ | v3 | **0.564** | 0.244 | 0.561 | 0.237 | 0.599 | 0.251 | 0.574 | 0.241 | 0.555 | 0.276 |
| $\text{MIX}_{\text{full}}$ | v3 | 0.837 | 0.236 | 0.421 | 0.179 | 0.424 | 0.259 | 0.585 | 0.316 | 0.702 | 0.255 |
| $\text{MIX}_{\text{minimal}}$ | v3 | 0.837 | 0.236 | 0.412 | 0.156 | **0.271** | 0.087 | **0.174** | 0.081 | **0.150** | 0.058 |

**Table S4 Mean and standard deviation of rRMSE results of all simulations and $n_{\textbf{metabolites}}$.**

| | Sim. | Mean | Std | Mean | Std | Mean | Std | Mean | Std | Mean | Std |
|---|---|---|---|---|---|---|---|---|---|---|---|
| $n_{\text{metabolites}}$ | - | 4 | | 10 | | 20 | | 40 | | 60 | |
| PQN | v1 | 0.410 | 0.024 | 0.165 | 0.037 | 0.083 | 0.015 | **0.054** | 0.009 | **0.047** | 0.009 |
| $\text{PKM}_{\text{full}}$ | v1 | **0.320** | 0.179 | **0.159** | 0.065 | 0.108 | 0.025 | 0.092 | 0.020 | 0.076 | 0.018 |
| $\text{PKM}_{\text{minimal}}$ | v1 | **0.320** | 0.179 | 0.283 | 0.168 | 0.279 | 0.163 | 0.310 | 0.192 | 0.278 | 0.153 |
| $\text{MIX}_{\text{full}}$ | v1 | 0.392 | 0.070 | 0.160 | 0.035 | 0.103 | 0.045 | 0.200 | 0.141 | 0.204 | 0.131 |
| $\text{MIX}_{\text{minimal}}$ | v1 | 0.392 | 0.070 | 0.160 | 0.036 | **0.082** | 0.015 | 0.057 | 0.010 | 0.049 | 0.010 |
| PQN | v2 | 0.409 | 0.026 | 0.269 | 0.045 | 0.173 | 0.043 | **0.107** | 0.022 | **0.082** | 0.019 |
| $\text{PKM}_{\text{full}}$ | v2 | **0.277** | 0.176 | **0.239** | 0.132 | 0.246 | 0.138 | 0.275 | 0.136 | 0.283 | 0.137 |
| $\text{PKM}_{\text{minimal}}$ | v2 | **0.277** | 0.176 | 0.283 | 0.168 | 0.300 | 0.164 | 0.299 | 0.175 | 0.289 | 0.162 |
| $\text{MIX}_{\text{full}}$ | v2 | 0.379 | 0.062 | 0.264 | 0.048 | 0.182 | 0.044 | 0.176 | 0.104 | 0.215 | 0.140 |
| $\text{MIX}_{\text{minimal}}$ | v2 | 0.379 | 0.062 | 0.254 | 0.047 | **0.166** | 0.039 | **0.107** | 0.022 | 0.083 | 0.019 |
| PQN | v3 | 0.408 | 0.026 | 0.218 | 0.044 | 0.130 | 0.024 | **0.081** | 0.014 | **0.064** | 0.013 |
| $\text{PKM}_{\text{full}}$ | v3 | **0.269** | 0.164 | **0.200** | 0.092 | 0.156 | 0.051 | 0.119 | 0.034 | 0.115 | 0.034 |
| $\text{PKM}_{\text{minimal}}$ | v3 | **0.269** | 0.164 | 0.283 | 0.168 | 0.296 | 0.169 | 0.298 | 0.163 | 0.291 | 0.186 |
| $\text{MIX}_{\text{full}}$ | v3 | 0.372 | 0.053 | 0.211 | 0.045 | 0.159 | 0.085 | 0.197 | 0.139 | 0.221 | 0.150 |
| $\text{MIX}_{\text{minimal}}$ | v3 | 0.372 | 0.053 | 0.209 | 0.044 | **0.128** | 0.024 | 0.083 | 0.016 | 0.066 | 0.014 |

**Table S5 $p$-values between RMSE measures for all tested normalization model combinations** for $n_{\text{metabolites}} = 60$ calculated with the two-sided pairwise non-parametric Wilcoxon signed-rank test.

| Simulation | | $\text{PKM}_{\text{minimal}}$ | $\text{PKM}_{\text{full}}$ | $\text{MIX}_{\text{minimal}}$ |
|---|---|---|---|---|
| v1 | $\text{PKM}_{\text{full}}$ | 5.8E−18 | | |
| v1 | $\text{MIX}_{\text{minimal}}$ | 3.9E−18 | 2.0E−13 | |
| v1 | $\text{MIX}_{\text{full}}$ | 1.9E−02 | 1.6E−17 | 4.0E−18 |
| v2 | $\text{PKM}_{\text{full}}$ | 1.0E−02 | | |
| v2 | $\text{MIX}_{\text{minimal}}$ | 4.3E−18 | 3.9E−18 | |
| v2 | $\text{MIX}_{\text{full}}$ | 3.2E−04 | 1.4E−01 | 5.0E−18 |
| v3 | $\text{PKM}_{\text{full}}$ | 5.8E−14 | | |
| v3 | $\text{MIX}_{\text{minimal}}$ | 5.3E−18 | 7.0E−15 | |
| v3 | $\text{MIX}_{\text{full}}$ | 1.6E−04 | 1.1E−16 | 8.3E−18 |

**Table S6 $p$-values between rRMSE measures for all tested normalization model combinations** for $n_{\text{metabolites}} = 60$ calculated with the two-sided pairwise non-parametric Wilcoxon signed-rank test.

| Simulation | | $\text{PKM}_{\text{minimal}}$ | $\text{PKM}_{\text{full}}$ | $\text{MIX}_{\text{minimal}}$ | $\text{MIX}_{\text{full}}$ |
|---|---|---|---|---|---|
| v1 | $\text{PKM}_{\text{full}}$ | 3.9E−18 | | | |
| v1 | $\text{MIX}_{\text{minimal}}$ | 3.9E−18 | 1.5E−16 | | |
| v1 | $\text{MIX}_{\text{full}}$ | 3.2E−04 | 1.1E−15 | 9.0E−18 | |
| v1 | PQN | 3.9E−18 | 1.7E−17 | 2.9E−03 | 5.8E−18 |
| v2 | $\text{PKM}_{\text{full}}$ | 8.5E−01 | | | |
| v2 | $\text{MIX}_{\text{minimal}}$ | 3.9E−18 | 3.9E−18 | | |
| v2 | $\text{MIX}_{\text{full}}$ | 1.2E−03 | 2.3E−06 | 1.1E−16 | |
| v2 | PQN | 3.9E−18 | 3.9E−18 | 5.9E−01 | 3.9E−17 |
| v3 | $\text{PKM}_{\text{full}}$ | 1.4E−16 | | | |
| v3 | $\text{MIX}_{\text{minimal}}$ | 3.9E−18 | 6.3E−18 | | |
| v3 | $\text{MIX}_{\text{full}}$ | 2.1E−03 | 6.3E−12 | 7.1E−18 | |
| v3 | PQN | 3.9E−18 | 5.0E−18 | 8.0E−02 | 6.1E−18 |

**Table S7 $p$-values testing significant reduction of RMSE and rRMSE in $\textbf{MIX}_{\textbf{minimal}}$ compared to all other tested normalization models** for $n_{\text{metabolites}} = 60$ calculated with the one-sided pairwise non-parametric Wilcoxon signed-rank test.

| | Simulation | $\text{PKM}_{\text{minimal}}$ | $\text{PKM}_{\text{full}}$ | $\text{MIX}_{\text{full}}$ | PQN |
|---|---|---|---|---|---|
| RMSE | v1 | 2.0E-18 | 1.0E-13 | 1.9E-18 | - |
| RMSE | v2 | 2.5E-18 | 1.9E-18 | 2.1E-18 | - |
| RMSE | v3 | 4.1E-18 | 3.5E-15 | 2.6E-18 | - |
| rRMSE | v1 | 4.5E-18 | 7.6E-17 | 1.9E-18 | 1.0E+00 |
| rRMSE | v2 | 5.5E-17 | 1.9E-18 | 1.9E-18 | 7.1E-01 |
| rRMSE | v3 | 3.6E-18 | 3.2E-18 | 1.9E-18 | 9.6E-01 |

**Table S8 Time spent in seconds for one optimization step of simulation v3** synthetic data with different $n_{\text{metabolites}}$ and normalization models.

| | Mean | Std | Mean | Std | Mean | Std | Mean | Std | Mean | Std |
|---|---|---|---|---|---|---|---|---|---|---|
| $n_{\text{metabolites}}$ | 4 | | 10 | | 20 | | 40 | | 60 | |
| PQN | 2.2E−04 | 4.7E−05 | 2.0E−04 | 3.1E−05 | 2.1E−04 | 3.4E−05 | 2.3E−04 | 3.4E−05 | 2.6E−04 | 5.7E−05 |
| $\text{PKM}_{\text{full}}$ | 1.6E+00 | 1.1E+00 | 4.1E+00 | 3.5E+00 | 9.4E+00 | 7.0E+00 | 3.7E+01 | 1.9E+01 | 1.1E+02 | 4.4E+01 |
| $\text{PKM}_{\text{minimal}}$ | 1.6E+00 | 1.1E+00 | 1.5E+00 | 1.1E+00 | 1.7E+00 | 7.4E−01 | 1.7E+00 | 6.2E−01 | 1.9E+00 | 1.2E+00 |
| $\text{MIX}_{\text{full}}$ | 1.9E+00 | 1.3E+00 | 4.6E+00 | 4.1E+00 | 4.5E+00 | 3.7E+00 | 1.1E+01 | 1.1E+01 | 1.9E+01 | 2.2E+01 |
| $\text{MIX}_{\text{minimal}}$ | 1.9E+00 | 1.3E+00 | 1.6E+00 | 9.8E−01 | 2.1E+00 | 1.4E+00 | 2.2E+00 | 1.3E+00 | 2.6E+00 | 2.3E+00 |

**Table S9 $p$-values of difference between rRMSE of $\text{MIX}_{\text{minimal}}$ and PQN** as tested with the one-sided, pairwise Wilcoxon signed-rank test for simulation v3 synthetic data for different noise fractions, $f_n$.

| $f_n$ | $p$-value |
|---|---|
| 0 | 9.8E−01 |
| 0.1 | 1.7E−08 |
| 0.2 | 3.0E−17 |
| 0.3 | 1.9E−18 |
| 0.4 | 1.9E−18 |
| 0.5 | 1.9E−18 |
| 0.6 | 1.9E−18 |
| 0.7 | 1.9E−18 |
| 0.8 | 2.4E−17 |
| 0.9 | 4.1E−10 |

### 3 Supplementary Equations

#### 3.1 Weighting the Loss Terms – Calculation of λ

$\lambda$ is an important hyperparameter in the MIX model optimization. It basically scales the loss functions of the PKM and PQN part of a MIX model relative to each other (Equations 9b, 9c). Here we show the derivation of the equation used to calculate $\lambda$.

$$\frac{\lambda}{(1-\lambda)} = \frac{1/n^{\text{PKM}}_{\text{data points}}}{1/n^{\text{PQN}}_{\text{data points}}} \tag{S1a}$$

where

$$n^{\text{PKM}}_{\text{data points}} = n^{\text{PKM}}_{\text{metabolites}}\ n_{\text{time points}} \tag{S1b}$$
$$n^{\text{PQN}}_{\text{data points}} = n_{\text{time points}} \tag{S1c}$$

thus

$$\frac{\lambda}{(1-\lambda)} = \frac{1}{n^{\text{PKM}}_{\text{metabolites}}} \tag{S1d}$$
$$\lambda = \frac{1}{(n^{\text{PKM}}_{\text{metabolites}}+1)} \tag{S1e}$$

where

$$n^{\text{PKM}}_{\text{metabolites}} = \ell. \tag{S1f}$$

#### 3.2 2-Metabolite Caffeine Subnetwork

Equations S2a and S2b show the analytical solution to the first order mass action kinetics derived from a reaction network as shown in the bottom of Figure 4.

$$\mathbf{c}^{\text{caffeine}}(t) = \frac{k'_1}{k'_3+k'_2-k'_1}\ [\exp(-k'_1\ t) - \exp(-(k'_2+k'_3)\ t)] \tag{S2a}$$
$$\mathbf{c}^{m}(t) = k'_1\ k'_2\ [G(k'_1,k'_4,k'_5) + G(k'_4,k'_1,k'_5) + G(k'_5,k'_1,k'_4)] + c^m_0\ \mathrm{e}^{-k'_4\ t} \tag{S2b}$$

with

$$G(x,y,z) = G(x,y,z,t) = \frac{\exp(-xt)}{(x-y)(x-z)}, \tag{S2c}$$
$$k'_5 = k'_2 + k'_3 \tag{S2d}$$

for

$$m \in \{\text{paraxanthine, theobromine, theophylline}\} \tag{S2e}$$

##### 3.3 Goodness of Fit Measures

$$\mathrm{RMSE}^{\mathrm{fit}} = \sqrt{\frac{1}{n_{\text{time points}}} \sum_{j=1}^{n_{\text{time points}}} \left(V_j^{\mathrm{true}} - V_j^{\mathrm{fit}}\right)^2} \tag{S3a}$$

$$\mathrm{rRMSE}^{\mathrm{fit}} = \sqrt{\frac{1}{n_{\text{time points}}} \sum_{j=1}^{n_{\text{time points}}} \left(O_j - \overline{O}\right)^2} \tag{S3b}$$

with

$$O_j = \frac{V_j^{\mathrm{true}}/V_j^{\mathrm{fit}}}{\mathrm{mean}\left\{\mathbf{V}^{\mathrm{true}}/\mathbf{V}^{\mathrm{fit}}\right\}} \tag{S3c}$$

for

$$j \in \{1, ..., n_{\text{time points}}\} \tag{S3d}$$

$$\mathrm{fit} \in \{\mathrm{PQN,\ PKM,\ MIX}\} \tag{S3e}$$
